# The HIV-1 Nuclear Export Complex Reveals the Role of RNA in Crm1 Cargo Recognition

**DOI:** 10.1101/2024.09.22.614349

**Authors:** Amber M. Smith, Yang Li, Arianna Velarde, Yifan Cheng, Alan D. Frankel

## Abstract

Crm1 is a highly conserved nuclear exportin that transports >1000 human proteins including ribonucleoprotein (RNP) complexes. The interface between Crm1 and RNP cargos is unknown. The HIV regulatory protein, Rev, was one of the first identified cargos for Crm1 and contains a prototypic nuclear export sequence (NES). We present the cryo-electron microscopy structure of the HIV-1 nuclear export complex (Crm1/Ran-GTP and the Rev/RRE RNP). Rev binds at a previously unseen protein-protein binding site that stabilizes a unique Crm1 dimer and positions two NESs within the Crm1 dimer. The orientation of Rev binding positions the RRE within a charged pocket on the inside of the Crm1 toroid, mediating direct RNA-Ran-GTP contacts, highlighting the significant role of the RRE in the interaction. Structure based mutations, combined with cell-based assays, show that Crm1 has multiple distinct cargo recognition sites and explains how Crm1 can recognize a diverse range of protein and RNP cargos.

## Introduction

Chromosome maintenance 1 (Crm1) is a member of the karyopherin-β (Kap) family, a group of proteins responsible for nucleocytoplasmic trafficking of proteins and RNA. Characteristic of the Kap family, Crm1 is made up of 21 tandem **h**untington, **e**longation factor 3, protein phosphatase 2**A** and **T**OR1 (HEAT) repeats that are organized into a toroid(1– 5). Currently, the number of Crm1 cargos exceeds 1000 human proteins and the functionalities of these proteins include ribosome biogenesis, mRNA degradation, autophagy, and cytokinesis(6). Crm1 cargos commonly contain a nuclear export sequence (NES)(6–11). The sequence and structural requirements for a Crm1 NES are loose(6, 7, 11–14) and the affinities of these NES peptides vary greatly from cargo to cargo(13–17). The variability in NES sequence and structure raises the question of how one protein can recognize and transport such a diverse range of proteins with different molecular characteristics. To add to the complexity of its cargo recognition, Crm1 is the only known karyopherin to export RNP cargos, in addition to proteins(18–21). Unlike other members of the Kap family, the stoichiometry of Crm1 also contributes to cargo recognition: the structure of Crm1 with Snurportin (SPN), revealed an interaction with a Crm1 monomer, whereas a low-resolution structure of the HIV-1 nuclear export complex (NEC), which uses the viral Rev protein and RNA to hijack host Crm1, revealed a Crm1 dimer(22, 23). In the SPN complex, which is the only high-resolution structure of Crm1 with cargo, SPN binds to the outer convex surface of the Crm1 toroid while the Ras-related nuclear protein (Ran), an essential Crm1 co-factor, binds to the inner concave surface(22, 24). The lack of interaction between SPN and Ran-GTP is an anomaly to Kap exportins, which couple cargo and Ran-GTP binding through direct contacts(25). Although the resolution is low, the NEC structure suggests a distinct mode of cargo recognition. These differences highlight the important outstanding question of how Crm1 interacts with such a large and diverse body of cargos.

Like all viruses, HIV requires host protein interactions to redirect critical host biological processes during replication. HIV relies on overlapping reading frames and alternative splice sites to translate all the necessary proteins for viral replication and infection. A single ∼9 kb viral transcript produced from the integrated viral DNA is fully processed (capped, spliced, cleaved and polyadenylated) by the host cell machinery early in the viral life cycle to yield ∼2 kb mRNA species encoding the Rev or Tat regulatory proteins or the Nef accessory protein(26, 27). Later in the life cycle, HIV bypasses the host mRNA processing machinery to facilitate the nuclear export of intron-containing transcripts, which encode the remaining viral proteins and genomic RNA. In the host cell, splicing and export of host mRNAs are tightly regulated in the nucleus, ensuring that intron-containing mRNAs are not exported to the cytoplasm(28– 30). For these host mRNAs, export proteins are recruited to the mature mRNAs via the spliceosome and are then shuttled to the cytoplasm by the protein TAP(31). In HIV, the Rev protein functions as an adapter to export its incompletely spliced and unspliced transcripts(32–34) by binding to the Rev response element (RRE) a structured region of the viral RNA located in a viral intron(35–37), and simultaneously to the Crm1 export receptor(38, 39). Rev is comprised of 116 amino acids and has three functional domains: a bipartite oligomerization domain (OD; residues 12-26 and 50-60), an arginine rich motif (ARM; residues 35-49) which functions as the RNA-binding domain and nuclear localization sequence (NLS), and a nuclear export sequence (NES; residues 75-83). Rev oligomerizes sequentially and cooperatively on the ∼351 nt RRE, binding first at the high-affinity RRE stem IIB site(40–47). The entire Rev/RRE RNP is comprised of six Rev subunits(48). Rev was one of the first proteins identified to contain an NES(7). As a component of a RNP and by utilizing a Crm1 dimer, Rev serves as an excellent model for elucidating some of the code to Crm1 cargo recognition.

In this study, we determined the structure of the HIV-1 NEC by single-particle cryo-electron microscopy (cryo-EM) to reveal how RNP cargos bind to Crm1, and how HIV hijacks Crm1 for its own replication. Additionally, we provide insights into the broader question of cargo recognition by Crm1. Our structure highlights the significance of the Rev-driven Crm1 dimerization and identifies two previously unseen cargo recognition sites (a protein-protein site and an RNA-protein site). These newly identified cargo binding sites led us to investigate how distinct each site is for particular cargos and whether some interactions might be individually targeted for therapeutics. These results emphasize how Crm1 can accommodate such an expansive repertoire of cargos with varied NESs by exploiting multipartite recognition and stoichiometric differences to generate functional export complexes.

## Results

### HIV-1 NEC Structure Description

Our early work demonstrated that Crm1 dimerizes when exporting the HIV Rev/RRE RNP from the nucleus to the cytoplasm, unlike monomeric Crm1 seen with other export cargos(23). Using this unique dimer formation as a criterion, we modified HIV-1 Rev (HXB3 isolate), HIV-1 RRE RNA (SF-2 isolate)(49), and human Ran to promote stable complex formation with a Crm1 dimer. Complexes were assembled with a Ran variant that combines two well-characterized modifications to prevent GTP hydrolysis(50–52), a Rev chimera with an engineered high-affinity NES (NS2Val)(13), and an RRE variant (RRE61)(49) that locks the RNA into a single conformation. In addition to the components of the NEC, we included a 113-amino acid FG-repeat polypeptide from the nuclear pore protein, Nup214 (residues 1916-2033), attached to the maltose binding protein (MBP) to increase Crm1’s affinity for the Rev/RRE RNP(53). Negative stain EM indicated that all modifications maintained a NEC structure indistinguishable from that with the wild-type HIV-1 Rev/RRE complex (Figure S1). Obtaining stable HIV-1 NEC allowed us to determine a structure with a global nominal resolution of 4.2 Å, with 3.7 Å local resolution for the Crm1/Ran-GTP dimer (Figures 1A-B, S2A and Table S1). The nominal resolution of the Rev/RRE portion is closer to 5.8-6.8 Å, consistent with an increase in conformational flexibility as a function of distance from the Crm1/Ran dimer (Figure S2B). An atomic model of the Crm1/Ran-GTP/Rev-RRE complex (Figure 1C) was reliably produced by rigid body docking of the crystal structures of Crm1/Ran-GTP(53) and Rev/RRE(54) followed by real space refinement(55, 56). The resolution of the Crm1/Ran-GTP dimer is sufficient to accurately trace the backbone and observe distinct helical pitch and β-sheets and some side chain densities. The residues that stabilize the Crm1 dimer are critical for Rev function and differ between primates and murines(57, 58). Although we modeled these residues at the Crm1 dimer interface in our previous 25 Å structure(23), the increased resolution of our current NEC structure (3.7 Å) confirms that these residues (V402, P411, M412, F414, R474, E478 and H481) in combination with a conserved residue (D471) create a highly polar interaction network flanked by hydrophobic residues that strengthen the human Crm1 dimer interface compared to its murine counterpart (Figures S3B and S4)(23, 57, 58). The resolution of the Rev/RRE RNP is adequate to accurately locate the α-helices of Rev and the stem IIB of the RRE. The architecture of the Rev dimer is driven by binding to the RRE, which sets the same Rev dimer crossing angle observed in the crystal structure of the Rev-RRE dimer complex alone(54) (Figure 2A). Connectivity between the NES and the OD (the OD/NES linker; ONL) is clearly observable for each subunit of Rev for the first time (Figure S3A). The density for both Rev NES peptides is clearly seen within the two Crm1 NES-binding clefts at the same resolution as the Crm1/Ran dimer.

**Fig. 1.**
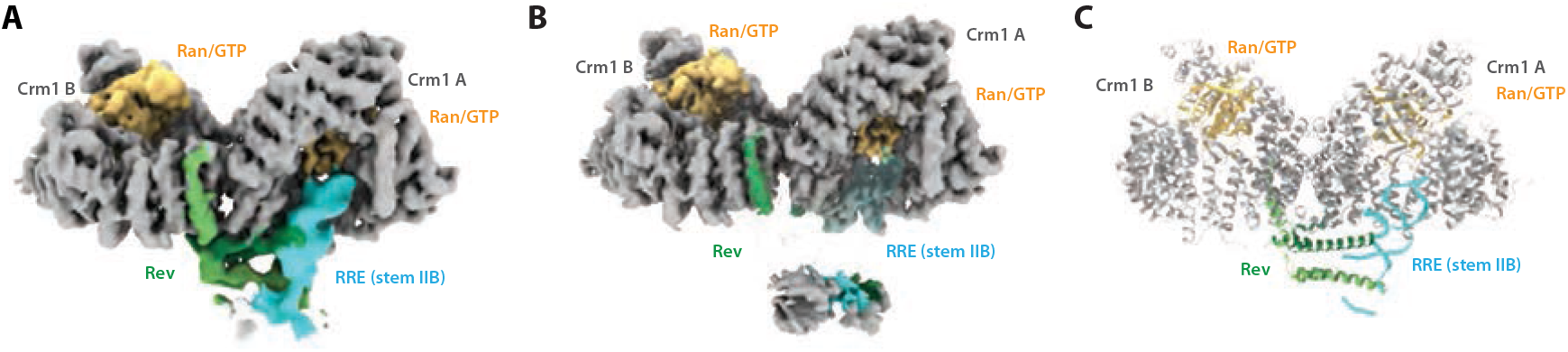
The HIV Nuclear Export Complex. (A) Cryo-EM density map of the HIV-1 nuclear export complex (NEC) colored by proteins. Crm1 in gray, Ran/GTP in peach, RRE (stem IIB) in cyan, Rev in light and dark green. (B) The sharpened cryo-EM map (DeepEMhancer) is colored by protein following the same color code in (A) with the locally filtered map shown in (A) as transparent. (C) Ribbon diagram of the HIV-1 NEC with the same coloring observed in (A).

**Fig. 2.**
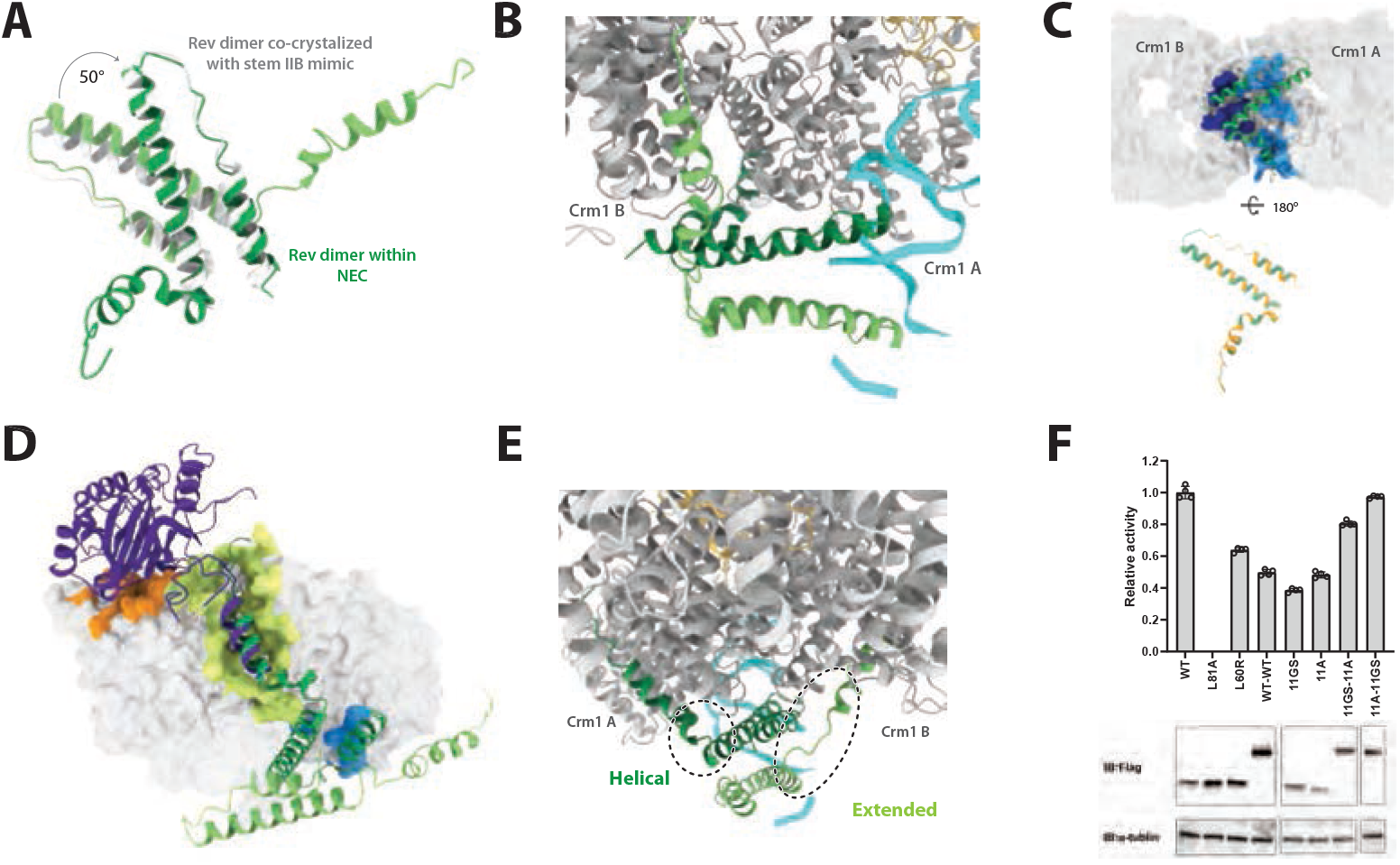
Rev dimer drives Crm1 dimer. (A) Overlay of the NEC Rev dimer (dark and light green) with the Rev dimer co-crystallized to a stem IIB mimic (gray; PDB: 4PMI). (B) Close-up view of the Rev dimer interacting with the Crm1/Ran-GTP dimer. Proteins/RNA are represented in ribbon diagram, Crm1 A and B in gray, Ran-GTP in peach, Rev dimer dark and light green, RRE in cyan. (C) Surface representation of the Crm1 dimer (transparent gray) and a ribbon diagram of the top Rev subunit (dark green). Residues on Crm1 that interact with Rev represented in light or dark blue. Residues on Rev that interact with Crm1 colored in orange. (D) Rev dimer (light and dark green) and SPN (PDB: 5DIS, purple) are shown in ribbon diagrams, with a monomer of Crm1 shown as a surface representation (transparent light gray). The residues on Crm1 that form the SPN binding site are colored in orange. The residues on Crm1 that form the Rev binding site are colored in blue. The residues of the NES binding cleft are shown in yellow. (E) Ribbon diagram of the side view of the NEC revealing the structural differences between the ONLs from each Rev subunit (dark green helical ONL; light green extended ONL). (F) Rev mediated nuclear export of RRE containing transcripts were assayed in HEK293T cells transfected with either wild type Rev, Rev L81A (an NES deficient mutant), Rev L60R, wild-type linked-dimer Rev, monomeric Rev ONL-11GS, monomeric Rev ONL-11A, linked-dimer Rev ONL-11GS/11A and linked-dimer Rev ONL-11A/11GS. These experiments were biologically replicated and normalized to wild type. Western blots below show protein expression for flag-tagged Rev and α-Tubulin as a loading control.

### Rev dimer drives the Crm1 dimer

At first glance, the Crm1/Ran dimer appears to be the dominant scaffold, but a closer examination reveals that a dimeric-Rev bound to stem IIB dictates the overall organization and stability of the entire complex. Previous structural and biochemical studies indicate that a Rev dimer is the most stable form of the protein and forms the core of the RNP(54, 59–63), and the NEC structure is consistent with this view. The Rev dimer binds to the Crm1 dimer with its dimer interface perpendicular to the dimer interface of Crm1 (Figure 2B). The upper subunit of the Rev dimer uses its NES to bind to the Crm1 subunit A and creates an extensive protein-protein interaction across the outer face of the Rev molecule, utilizing the OD, and interacting with both Crm1 subunits at a novel protein interaction site on the outside of the Crm1 toroid (Figures 2C and 2D). The NES of the lower Rev subunit binds the Crm1 subunit B without any additional interactions. Dimerization of Crm1 and Rev doubles the total buried surface area and is most likely the reason why nuclear export is achievable with Rev’s weak NES(16). The structure of a Crm1-SPN complex, which is monomeric, similarly shows interactions via its NES and a protein binding site on the outside of the Crm1 toroid(22, 24, 53). The SPN and Rev binding sites, located on the outside of Crm1, contrast with all other Kap family members which bind their cargos on the inside of their HEAT repeats(3, 4, 64–74).

The C-terminal portion of Rev (residues 70-116, which includes the NES) has not been observed as a defined structure prior to this work. Low-resolution density of this portion was observed using cryo-EM of Rev filaments with the protein alone, suggesting that it is partially ordered in the filament context(60). In the NEC structure, this region becomes completely ordered when Rev is bound to Crm1, and we can observe clear density past the end of the NES (residue 90). Interestingly, the asymmetry of the complex is apparent not only in the orientation of the Rev/RRE but also within the Rev dimer, as suggested by previous biochemical studies(48). Each Rev NES interacts with its corresponding Crm1 NES cleft in the same well-characterized manner(13, 22) but the stacking arrangement of the Rev dimer results in differences in distance between each Rev subunit to the NES clefts of Crm1. As a consequence, the adjacent linkers in Rev (ONLs) adopt different conformations (Figure 2E). The upper Rev ONL is helical whereas the lower Rev ONL adopts an extended confirmation, consistent with previous experiments showing that substituting the linker with alanines or with a Gly-Ser repeat, which affects the helical propensities of both Rev subunits, reduced nuclear export and replication activities(75). To validate this asymmetry, we generated linked Rev dimers in which the N-terminal Rev molecule contained a helical 11-alanine (11A) ONL and the C-terminal Rev a flexible 11-Gly-Ser (11GS) ONL, or the converse orientation. Both linked dimers showed increased nuclear export activity compared to monomeric Rev containing either the 11A or 11GS substitutions (Figure 2F), and notably the dimer containing the N-terminal Rev with high helical propensity of the ONL (11A) and a C-terminal Rev with extended/flexible propensity of the ONL (11GS) was as active as wild-type Rev. In contrast, a linked dimer with wild-type ONLs exhibited low export activity, as expected given that Rev does not tolerate additions to its N- or C-termini(75). These results authenticate that the orientation of the Rev dimer in our model is correct.

### Rev oligomer stabilizes the Crm1 dimer

The bipartite OD is comprised mostly of hydrophobic amino acids which create both the Rev dimer interface and a higher-order oligomer interface using opposite sides of the OD helices(48, 54, 59–62, 76). In the absence of Crm1, the Rev/RRE RNP forms discrete hexameric complexes(48). In the presence of Crm1, molecular weight characterization experiments confirm that six molecules of Rev are also bound to the RRE(23). Studies have identified an oligomer deficient Rev mutant (L60R) that does not support virus replication(77) and prevents the formation of discrete Rev/RRE RNP complexes observed with wild-type Rev(40, 42, 48, 76). It is the OD of Rev in the NEC structure that contributes to the protein-protein binding site on Crm1, illustrating why mutations in this region are detrimental to complex formation. We do not observe the additional four Rev subunits in the cryo-EM structure due to inherent flexibility of the longer RNA and its bound proteins, however it is clear biochemically that six subunits are bound and that the complex cannot be formed without the full repertoire of RNA and six Rev subunits. To investigate how the Rev oligomer contributes to NEC stability, we examined complexes assembled with an oligomer deficient Rev mutant (L60R). Negative stain 2D class averages with L60R exhibited a mixture of monomeric and dimeric forms of Crm1, unlike the fully dimeric complexes observed with wild-type Rev (Figure S5). Interestingly, nuclear export activities of either L60R or an L60A mutant in the context of Rev from the NL4-3 viral isolate resulted in 30% and 80% activity(40, 77) yet neither virus replicates, indicating a high threshold of export activity needed for replication(77). We observed a similar decrease of L60R activity in HXB3 Rev used in the NEC structure as well as for the linked wildtype Rev dimer (Figure 2F), correlating with their mixtures of Crm1 monomers and dimers (Figure S5). While we can confidently place a single Rev dimer in the NEC, the additional Rev subunits are likely bound to the extended and flexible RRE and thus not observed in the current structure. These data reveal that a Rev dimer bound to stem IIB is largely sufficient to stabilize the Crm1/Ran dimer but that other portions of the Rev/RRE complex enhance NEC stability to meet the threshold for viral replication(78).

### Protein-Protein Binding Site is not Critical for RNP Cargo Recognition

Previous Crm1 structures, including Crm1-SPN, show a disordered loop between HEAT repeats 8 and 9(22, 53), whereas in the NEC structure, loop residues 389-400 in both Crm1 subunits become ordered (Figure 3A and S3C) and point towards the Crm1 dimer interface where Rev binds to fasten the Crm1 dimer (Figure 3B). The charge of the electrostatic surface of the Crm1 dimer in this region diminishes when the loops are ordered, instead creating a hydrophobic platform for Rev binding. This ordered loop is separated from an acidic loop by helix A of HEAT 9. The acidic loop is conserved among members of the Kap family (Figure S6)(1, 24) and plays a role in Ran-GTP and cargo recognition(22, 24, 79). In an apo-Crm1 structure (without Ran or cargo)(79) the C-terminal helix (helix B of HEAT 21) extends diagonally across Crm1 extending the overall Crm1 structure into a super helix that moves HEAT 11 and HEAT 12 toward each other, closing the NES cleft and preventing cargo from binding. The acidic loop folds back onto HEAT 10, HEAT 11 and HEAT 12, causing a rearrangement in the side chains that enables HEAT 11 to move towards HEAT 12. The acidic loop is further stabilized by a salt bridge to the C-terminal helix. When Ran-GTP is bound to Crm1, the C-terminal helix and the acidic loop undergo a conformational switch(24, 53, 79). The C-terminal helix moves back into a parallel orientation to helix A of HEAT 21 and interacts with HEAT 2 to HEAT 5, closing Crm1 into a toroidal shape structure and opening the NES cleft. The acidic loop reaches across the Crm1 toroid and binds to the basic region on Ran that occupies the central cavity (Figure S6)(24, 53). These conformational changes were the accepted explanation for why direct interactions between cargo and Ran-GTP were not required as in the case of the Crm1-SPN complex. The proximity of the ordered 389-400 loop to the acidic loop couples Rev/RRE and Ran binding even though Rev binds to the outside of the Crm1 toroid. The ordering of these loops to generate the Rev binding site infers that the Rev binding site is a critical interface and that the Crm1 cargo recognition for the Rev/RRE RNP follows the same rules observed in the Crm1-SPN complex but at a different binding site.

**Fig. 3.**
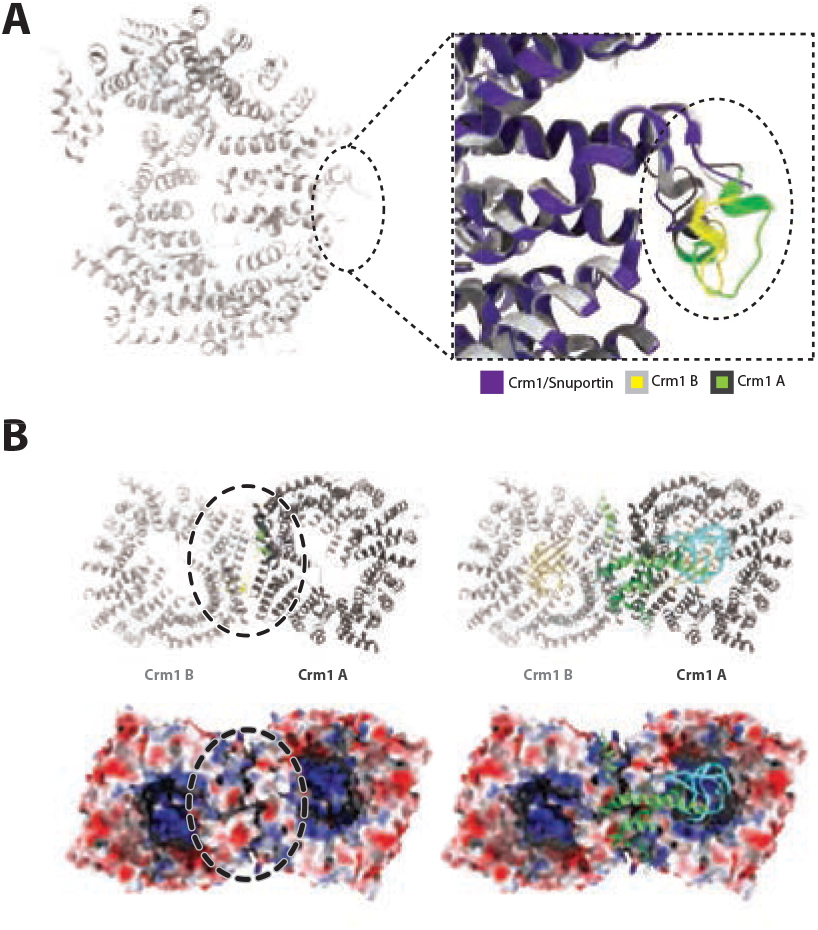
Disordered to ordered transition of Crm1 loops generate Rev binding site. (A) Binding of the HIV-1 RNP induces an ordering of a loop (residues 389-400) that is disordered in previously solved crystal structures. Overlay of Crm1/SPN crystal structure (purple, PDB: 5DIS), Crm1 A (dark gray/loop shown in green), and Crm1 B (light gray/loop shown in yellow). (B) The ordered loops are part of the Crm1 dimer interface. Top Left: ribbon diagram of the Crm1 dimer (light/dark gray) with the loops colored in yellow and green. Top Right: a ribbon diagram of the NEC, including Ran-GTP (peach) and the Rev-RRE RNP (light and dark green/cyan) on top of the Crm1 dimer shown to the left. Bottom Left: surface representation of the electrostatics of the Crm1 dimer with the Rev binding site highlighted with a dashed oval. Bottom Right: a ribbon diagram of the Rev-RRE RNP (light and dark green/cyan) on top of the electrostatic surface representation to the left.

The presence of multiple distinct binding sites would explain how Crm1 recognizes and interacts with such a large inventory of cargos. To test if these newly discovered sites and the SPN site are indeed segregated, each was mutated by changing all non-hydrophobic residues to alanines in various combinations (Figure S7). The mutational tolerance of Crm1 has not been systematically examined, although mutations to the NES binding cleft have been reported but none around the outside of the Crm1 toroid(5, 11, 80). Every Crm1 variant tested pulled down Ran from cells to a similar extent (data not shown), and each was analyzed by negative stain EM to ensure that the mutations did not affect the overall structure of Crm1 (Figure S8). Mutating 13 residues of the SPN binding site significantly impaired SPN binding (M1; Figure 4A), while reducing the number of mutations sequentially restored SPN binding, with two single-point mutations fully restoring the Crm1 interaction (M6 and M7; Figure 4A). Unexpectedly, mutations in the Rev binding site actually enhanced SPN binding in most cases compared to wild-type Crm1, and similarly, mutations in the SPN binding site enhanced Revmediated nuclear export activity (Figure 4B). Interestingly, mutating the Rev protein binding site in Crm1 had little effect on Rev activity, implying the RNA-protein interface contributes more to the Rev/RRE RNP interaction to Crm1 than the protein-protein interface and demonstrates the diversity in Crm1 multipartite cargo recognition.

**Fig. 4.**
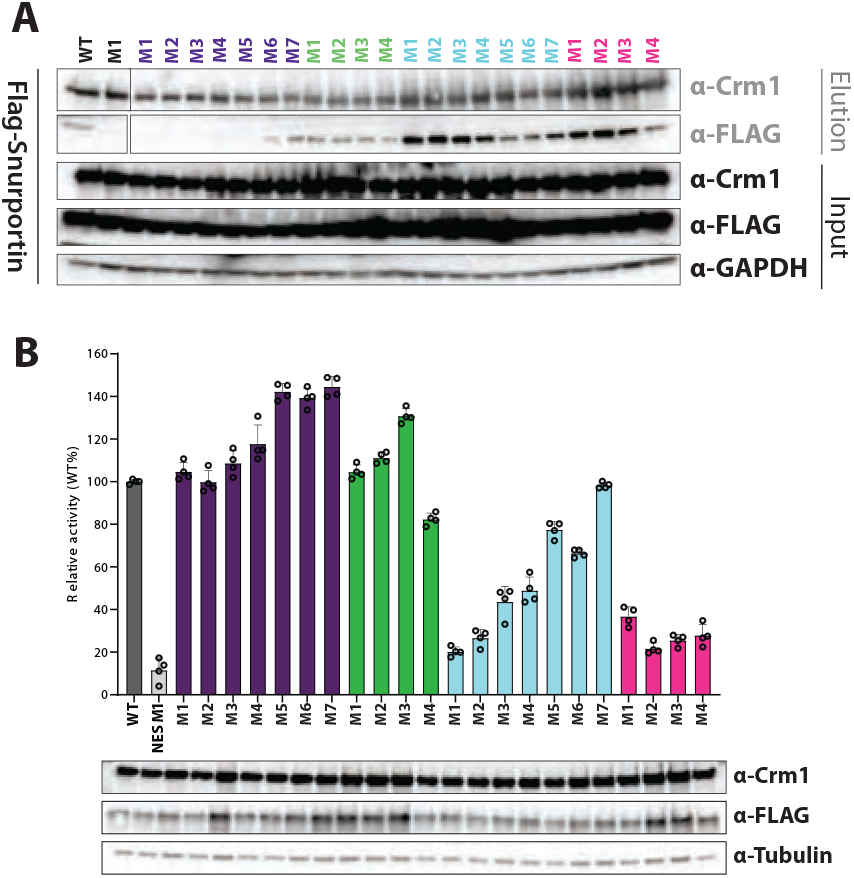
Crm1 Binding Sites are Separate and Distinct. (A) Co-IP analysis (2x Strep-Tag Pull-down) and Western blot analysis (anti-Flag and anti-Crm1) of Expi293 cells transfected with Crm1 mutant plasmids encoding Strep-Crm1 and Flag-Snuportin for 48 hours. Anti-GAPDH was used as a loading control. Residues altered in each mutant are shown in Figure S9. (B) Rev mediated nuclear export of RRE containing transcripts were assayed in HEK293T cells transfected with either wild-type Crm1 or Crm1 with mutations made to either the SPN, Rev, RRE or Rev/RRE binding sites. These experiments were biologically replicated and normalized to wild-type activity. Western blots below show protein expression for flag-tagged Rev and α-Tubulin as a loading control.

### RNA binding site is a Critical Interface

The NEC structure reveals an unexpected RNA-protein interface between Crm1/Ran and RRE stem IIB, in addition to the Rev-RNA interaction. The crystal structure of the dimeric Rev/RRE stem IIB complex(54) permitted us to model the RNA into the NEC map, confidently positioning the RNA helix and Rev dimer. Neither the open nor closed conformations observed in the recent crystal structure of stem loop II utilizing a tRNA-scaffold matches the conformation we observe in our structure(81). The IIB stem of the intact RRE is longer than that used in the co-crystal structure(54) and is precisely the length that fits into the NEC reconstruction. The electrostatics of Crm1 and Ran show a positively charged pocket where the RRE is cradled between Rev and Crm1/Ran (Figure 5A). Ran contains two loops (91-95, 122-135; Figure S9) that create a positively charged cap in the center of the Crm1 toroid that effectively limits the length of the RNA and positions the Rev dimer to bind the Crm1 dimer. Three residues in Ran (Arg95, Lys132 and Lys134) show clear density connected to the RRE and are the same residues that interact with the RNA cargos in the Expo-5 and XpoT structures(3, 4) (Figure 5B). The electrostatics of Crm1 reveal an asymmetric positively charged patch that extends from the Rev dimer across Ran to the other side of Crm1 (Figures 5A and 5B) and illustrates how Rev mimics a highly conserved RNA-binding mode within this family of trafficking proteins(3, 4, 82) (Figure 5C). XpoT and Expo-5 transport tRNA or miRNA in similar manners by wrapping around the RNA but neither requires adaptor proteins(83–86). RNA-interacting residues in the NEC are found within some of the HEAT repeats and linkers (Table S2) and are positioned within the composite Crm1/Ran site to match interactions between the Rev dimer, the Crm1 dimer and the RRE.

**Fig. 5.**
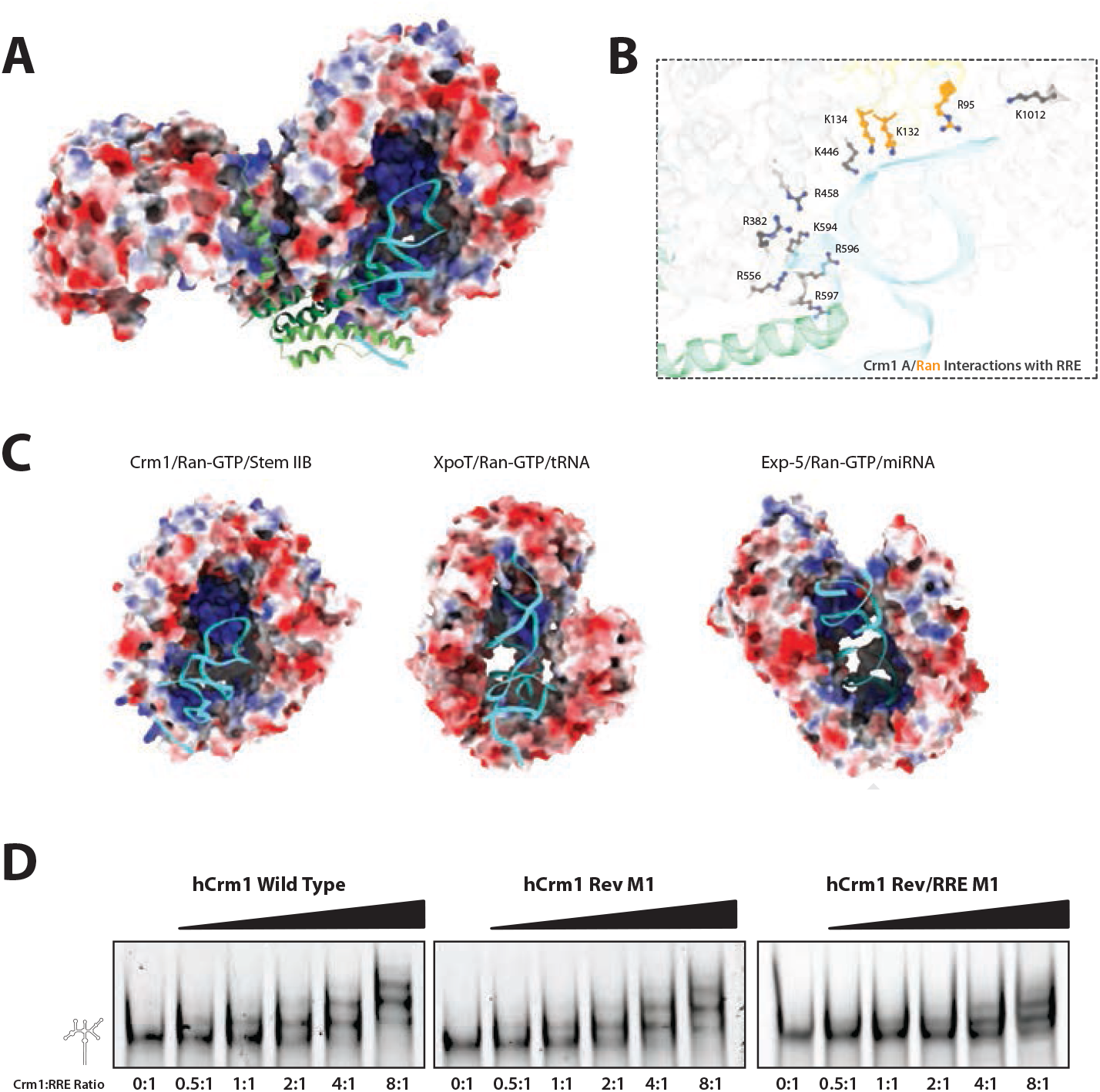
Rev hijacks a native RNA binding mechanism utilized by RNA trafficking exportins. (A) The NEC with the Crm1/Ran-GTP dimer shown in a surface representation colored by electrostatics with the Rev-RRE RNP shown in a ribbon diagram (Rev dimer in light and dark green, RRE in cyan). (B) Close-up view of the residues from Ran-GTP (orange) and Crm1 (dark gray) that interact with stem IIB of the RRE. (C) Electrostatics of each RNA trafficking exportin (left to right: Crm1, XpoT; PDB: 3ICQ and Exportin-5; PDB: 3A6P) with Ran-GTP shown in a surface representation colored by electrostatics, RNA cargos are shown in ribbon diagrams in cyan. (D) Wild type Crm1, Crm1 Rev M1 and Crm1 RevRRE M1 were combined at 6 different protein concentrations (0-800 nM) with HIV-1 RRE 234 (100 nM) and the complexes formed were analyzed by electrophoresis on a nondenaturing polyacrylamide gel.

To test if disrupting the RRE binding site affects Rev-mediated nuclear export activity, we generated Crm1 mutations at positions inside the toroid. Mutating 8 residues significantly decreased Rev activity, to a level similar to that of a triple mutant in the NES cleft (NES M1; L525A, K568A and F572A), emphasizing the significance of the RNA interaction in the NEC (Figure 4B). Reducing the number of mutations sequentially restored Rev nuclear export activity. As expected, mutating both the Rev and the RRE binding sites simultaneously decreased Rev activity more than mutating either alone. These results mirror those observed for the SPN-Crm1 interaction when mutations were made to the SPN binding site, indicating that RRE binding is as important to Crm1 recognition of the Rev/RRE RNP as SPN binding is to the Crm1 recognition of SPN. From these results, we may anticipate that protein and RNP cargos will show quite distinct rules of engagement with Crm1.

To further test the significance of the RNA-Crm1 interaction, we examined whether Crm1 might bind directly to the RRE even in the absence of Rev (Figure 5D). Wild-type Crm1, as well as a Rev binding site mutant, similarly bound to the RRE, forming higher-order multimeric complexes. The RNA became almost fully bound at a 4-fold excess of Crm1 over RRE. In contrast, a significant amount of unbound RNA remained at an 8-fold excess of a Crm1 mutant containing mutations in both the Rev and RRE binding sites. Negative stain EM (Figure S8) supported the biochemical results for all three bindings sites, as judged by Crm1 dimer formation, and together reveal the priorities of cargo recognition for the assembly of the NEC. NES M1 completely abolished Crm1 dimer formation (Figure S8), consistent with the importance of the NES for Rev-mediated nuclear export activity(87) and indicating that the Rev and RRE-Crm1 interactions are insufficient without NES binding. NES M1 also abolished SPN binding further establishing the essential role of the NES. Mutations to the Rev and RRE binding sites, either individually or together, significantly reduced but did not completely abolish dimer formation, highlighting the dominant effect of two NES peptides (Rev M1, RRE M1, Rev/RRE M1; Figure S8). Mutations to the SPN binding site prevented SPN binding to Crm1, while mutations to either the Rev or RRE binding sites did not alter the Crm1-SPN complex. Taken together with our biochemical experiment, it appears that the NES and protein-protein binding site are equally relevant in the case of SPN. These results reinforce the importance of two NES peptides of the Rev dimer binding to Crm1, as well as Crm1-RRE binding. Additionally, these results elucidate the differences between monomer vs dimer and protein vs RNP cargos.

### Identification of New Crm1 Recognition Sites

To further explore the diversity of Crm1 binding sites for cargo recognition, we examined the impact of Crm1 mutants on Survivin, another Crm1 cargo. Survivin is a member of the chromosome passenger complex (CPC) along with the proteins Aurora B, Borealin, and INCENP (inner centromere protein)(88). The CPC is a highly conserved complex that is responsible for the movement and segregation of chromosomes during meiosis and mitosis(89–94). The ability of the CPC to target the centromeres of chromosomes is dependent on the presence of Crm1 and the NES of Survivin(95). In fact, Crm1 was first identified in yeast as a temperature-sensitive mutant that caused disorganization of chromosomes, hence the name chromosome maintenance 1(96). To assess whether the Survivin cargo utilizes any of the three Crm1 binding sites identified in this current study, we tested the ability of Crm1 variants with each binding site fully mutated (SPN M1, Rev M1, RRE M1 and Rev/RRE M1) to pull down Survivin. Remarkably, every variant pulled down Survivin (Figure 6), even including a control with the NES cleft mutated (NES M1). These results suggest that Survivin binds to a fourth site on Crm1 yet to be identified (Figure 7). As mentioned, Survivin is part of the CPC, which also includes Aurora B known to interact with Crm1(6), and thus Survivin binding to the Crm1 NES cleft mutant may be mediated via Aurora B. Identifying the Survivin binding site on Crm1 and how the CPC interfaces with Crm1 will require further investigation.

**Fig. 6.**
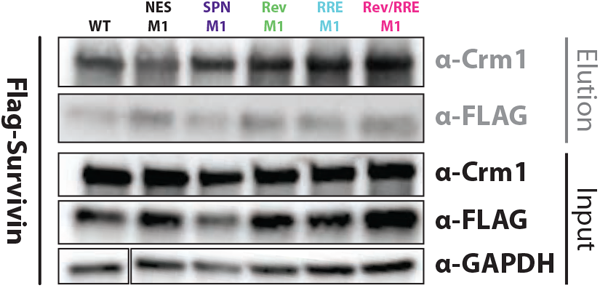
Survivin Binds to a Fourth Crm1 Binding Site. Co-IP analysis (2x Strep-Tag Pull-down) and western blot analysis (anti-Flag and anti-Crm1) of Expi293 cells transfected with plasmids encoding Strep-Crm1 and Flag-Survivin for 48 hours. Anti-GAPDH was used as a loading control.

**Fig. 7.**
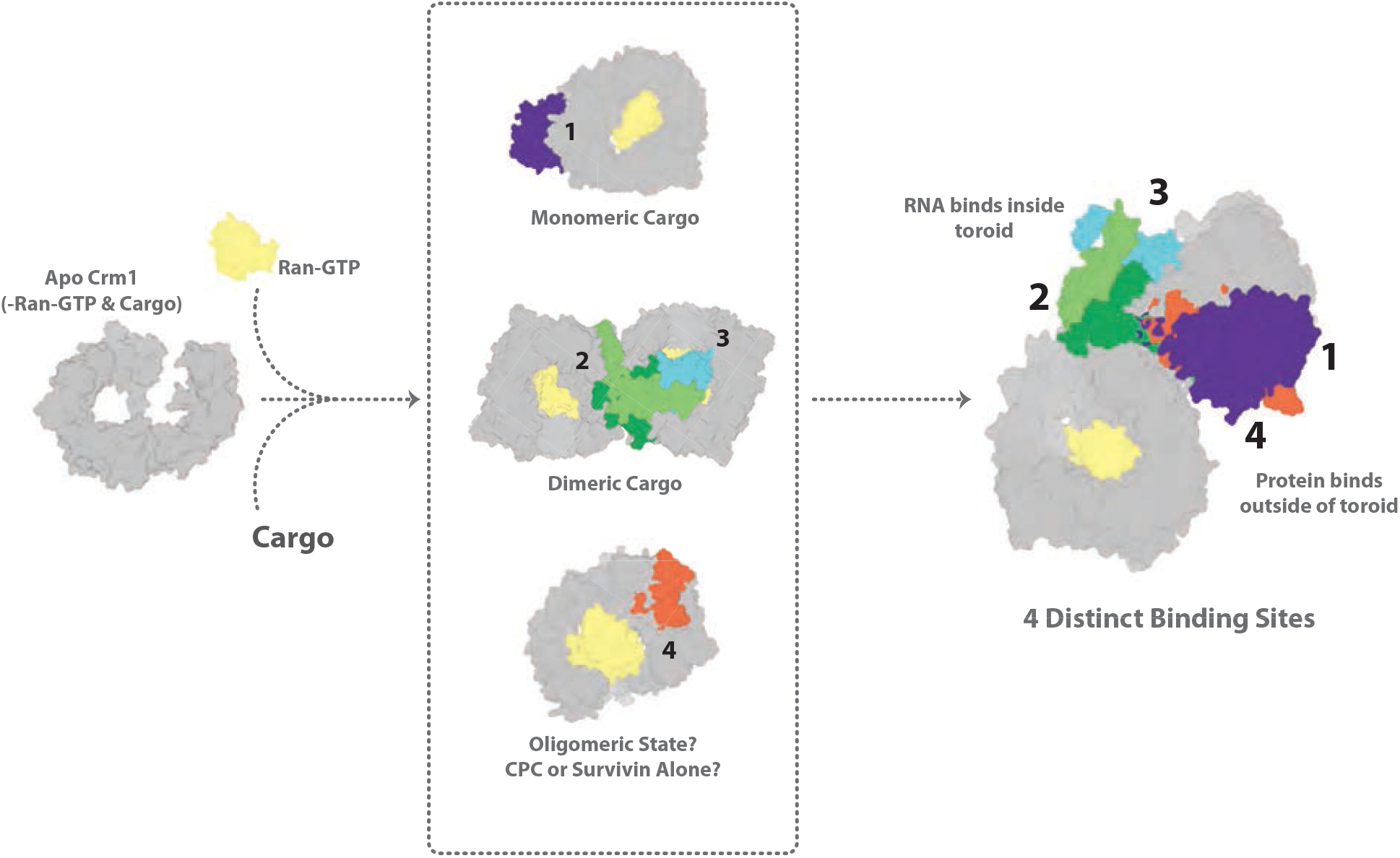
Crm1 Cargo Recognition Model. Left to Right: Crm1 in its apo form adopts a conformation that does not support Ran or cargo binding. In the presence of Ran-GTP, Crm1 interacts with its cargo based on cargo stoichiometry (Top: monomer for SPN, PDB:5DIS, Middle: dimer for Rev, Bottom: modeled schematic for Survivin, the oligomeric state or binding site is unknown but biochemical studies infer it is not the same as SPN or Rev). The current structures of Crm1 identify three separate and distinct cargo recognition sites (1, 2, 3). Pull down experiments between Crm1 and Survivin infer that there is a fourth binding site for Survivin. Crm1 exhibits a hybrid binding mode, where proteins bind on the outside of the Crm1 toroid and RNA binds to the inside of the toroid.

## Discussion

### Decoding Crm1 Cargo Recognition

Crm1 is a critical exportin that plays roles in many key biological processes. Understanding how this single protein is responsible for the trafficking of such a large repertoire of proteins and RNP complexes has long been a subject of inquiry. Crm1 is unique among the Kap family, for several reasons: it has expansive cargo diversity covering broad and unrelated biological functions; it utilizes a distinct mode of cargo interaction where cargo binds to the outside of the HEAT repeats without direct interactions to Ran-GTP; and it has the ability to traffic RNP complexes in addition to protein cargos. With only one structure of a Crm1-cargo complex (Crm1-SPN)(22, 24, 53) solved to date, it has not been possible to comprehensively understand how this single exportin can recognize such an extraordinary range of cargos that play such important and diverse roles in the cell. HIV Rev was one of the first proteins identified to contain an NES and to interact with Crm1(7, 39), and various Rev complexes and parts of the NEC have been investigated through numerous structural and biochemical studies over three decades. The structure of the entire NEC reported here caps efforts to understand how Rev facilitates the nuclear export of intron containing HIV transcripts while also beginning to establish a code for Crm1 cargo recognition.

Crm1 cargo recognition has traditionally focused on the NES, all efforts to modulate Crm1 activity has aimed at inhibiting the Crm1-NES interaction. However, this narrow approach overlooks other aspects of the protein/RNP complex that contribute to the interaction and fails to provide meaningful details. Our findings establish a new paradigm in Crm1 cargo recognition, showing that the second aspect of multipartite interactions is just as critical as the NES in ensuring the proper export of cargo. The Crm1-RRE interaction ensures that Rev is not exported until it is bound to the RRE. Structures of Rev have shown that dimer formation occurs independently of the RRE(59–62). Separating Crm1 recognition between Rev and the RRE ensures the intact Rev/RRE RNP is exported.

This concept aligns with Crm1’s recognition of SPN, where multipartite interactions also play a key role in directing proper export. Crm1’s interaction with the m3G cap binding domain of SPN regulates its affinity to Crm1, ensuring that the correct SPN state is exported(16, 24). SPN is an adaptor protein for the import of U small nuclear RNPs (U snRNPs), the Crm1-m3G cap interaction prevents Crm1 from exporting SPN until after it has released the U snRNPs in the nucleus(16). Examining the Crm1-NES interaction alone would not reveal the insights gained from analyzing both facets of binding. Each Crm1 cargo possesses distinct molecular characteristics, it is unrealistic to expect that a single surface could accommodate them all. Additionally, targeting the NES alone may be insufficient for some cargos, as in the case with Survivin that demonstrated Crm1 binding even when the NES cleft is mutated (NES M1; L525A, K568A and F572A). By comparing just two structures, we have already begun identifying features of Crm1 that deviate from the conventional understanding. Our work addresses how Crm1 accommodates a diverse array of cargos and introduces a new conceptual approach for studying each Crm1 cargo. This broader perspective offers a way to target specific cargos without disrupting Crm1’s essential functions. Investigating these alternative interactions opens new avenues for modulating nuclear transport and may reveal previously unknown aspects of Crm1 function.

### RNA is Integral to Cargo Recognition

The NEC structure is remarkable in that it exposes a rather extensive interface between Crm1/Ran and the HIV-1 Rev/RRE RNP involving many protein-protein and protein-RNA surfaces. The direct interactions between the RNA and Crm1/Ran reveal an intimate and extensive role for the RRE that is supported by biochemical and structural experiments. Previous studies utilizing a trans-dominant negative form of Rev (RevM10), a variant containing two mutations in the NES, uncovered an RRE mutant that rescued Rev activity by structurally altering the RRE(49, 97) and hinting at a connection between the NES and RNA binding. Our studies reveal, that like SPN, the Rev/RRE RNP utilizes multipartite cargo recognition. Both shared the requirement for the NES but unlike SPN, the Rev/RRE RNP depends on the RNA-protein interaction with Crm1 and not the Rev-Crm1 protein-protein interaction. The direct RNA interaction with Crm1/Ran observed in the structure helps to explain specific features of the Rev/RRE complex and may be relevant to other RNA cargos. Rev binding initiates at stem IIB of the ∼351 nt RRE with picomolar affinity(40, 46, 47, 98–101) and extends to a second weaker site in stem IA(40, 42, 102) and a third in stem I(40, 41, 43– 47). Based on the NEC structure, it is clear that the spacing between the bound Rev dimer and the RNA interacting loops of Ran-GTP could not accommodate an RNA much larger than IIB, making it clear why the Rev/RRE dimer forms the tight core of the NEC. The lack of structural information for the remainder of the Rev/RRE RNP likely reflects the intrinsic flexibility of RNA and the relatively less specific binding of Rev subunits to the other identified RRE binding sites(40, 42, 101). The asymmetric binding of the RRE to one molecule of the Crm1/Ran dimer, despite identical RNA binding pockets in each subunit, further illustrates the importance of Rev in driving Crm1 dimer formation. It uses a binding mode for RNA conserved among Kap exportins (Figure 5C)(3, 4) and further increases the buried surface area of the Rev/RRE cargo through extensive protein-protein surfaces, and dimerization of both cargo and exportin. The electrostatic nature of the composite Crm1/Ran RNA binding site and its conservation to other RNA exportins, Expo-5(4) and Xpo-T(3) (Figure 5C), suggest related binding modes for other RNP cargos and is yet another demonstration of the ability of a virus to mimic host interactions. Crm1 is the only exportin known so far to export RNPs(25). It appears to be a remarkably adaptable Kap in which its hybrid binding mode allows it to interact with both components of an RNP cargo, with protein bound on the outside of the toroid and RNA on the inside. The necessity for the NES explains why Crm1 does not traffic RNA alone.

### The Additive Impact of Two NES Peptides

Our studies demonstrate how the dimeric nature of Rev, specifically the presence of two NES peptides, drives and supports the dimeric form of Crm1. The Crm1 dimer in the NEC utilizes evolved species-specific residues within the Crm1 dimer interface(23, 57, 58) to form a docking site for the Rev/RRE RNP. It is not yet known if Crm1 dimerization is important in other cargo interactions, either for viral or host cargos but the evolution of these residues in humans and primates and their role in HIV infection suggests that the Crm1 dimer is important for host complexes as well. The core Rev dimer binds across the Crm1 dimer interface and utilizes hydrophobic residues from the higher-order oligomerization surface of Rev to further fasten the Crm1 dimer. This is an unexpected role for the OD and explains the reduced Rev activity and incomplete complex formation observed with mutants in this interface(40, 76, 77). The Rev dimer forms an identical subunit crossing angle in both the co-crystal structure of Rev/RRE(54) and in the NEC structure which, combined with the ability of the Rev ONL to adopt different ordered conformations, allows the two NES peptides from a single Rev dimer to grasp the two NES binding clefts in the Crm1 dimer. The presence of the two NES peptides appears more critical to NEC assembly than the Rev or RRE binding sites on Crm1. Mutating both the Rev and RRE binding sites simultaneously still allows a small population of the NEC to form whereas mutations to the NES cleft completely abolishes NEC assembly. In comparison to the Crm1-SPN complex, which is monomeric, the single SPN NES cannot compensate for the loss of the SPN binding site. These finding begin to establish rules that govern monomeric verse dimeric cargos. For the monomeric SPN, the protein-protein binding site and the NES binding site are equally important, as no complex was detected when either site was mutated. Unlike the SPN case, the NES binding site in the Rev/RRE RNP complex appears far more important than the Rev protein binding site because two NES are bound. Nonetheless, the protein-protein binding site between Crm1 and Rev orients the Rev/RRE RNP so that the two NES peptides and the RRE stem IIB are positioned correctly in the NES clefts of Crm1.

### Multiple Binding Sites Diversify Cargo Recognition

The extensive types of interactions observed in the NEC structure emphasize the value of multiple binding surfaces that extend well beyond the NES interaction. These multiple interfaces allow resilience to disruptions to one surface without eliminating the nuclear export ability of Crm1. In essence, Crm1 displays many possible surfaces to capture different cargos, which is further emphasized by Survivin. Survivin binding was permitted when all three known binding sites were mutated, as well as the NES cleft, raising the possibility that Crm1 interacts with other proteins of the CPC heterooligomer to compensate for the loss of the NES cleft. Thus, multipartite recognition and compensating interactions will likely further expand the modes in which tight binding are possible and thereby expand an already long list of Crm1 cargos(6, 103–106). The mutational plasticity of Crm1, the segregation of the SPN and Rev/RRE binding sites, and the discovery of a potential fourth binding site for Survivin make it reasonable to anticipate that other sites around the toroid for cargos with different molecular characteristics will be discovered. In the case of Rev, the binding site in part spans the Crm1 dimer interface and involves the dimeric Rev/RRE cargo, but the site might also be available to monomeric cargos. The dexterity of Crm1 to dimerize and adapt to and be stabilized by the oligomeric Rev complex suggests that other Crm1 cargos may also have oligomeric states (Figure 7) and indeed, recognition could extend to even higher order oligomeric complexes. The versatility of Crm1-cargo recognition extends in multiple directions.

These strategies highlight the plasticity with which Crm1 recognizes diverse cargos with a wide range of NES affinities(13–17). The ability to catalog Crm1 cargos by their recognition modes may make it possible to target specific groups of cargos therapeutically based on their interactions. The majority of the identified Crm1 cargos are proteins that have a very strong bias for the cytoplasm(6). Current models of the nuclear pore complex (NPC) suggest that it is far more permeable than previously believed and does not have a clear size cutoff(107, 108). Thus, Crm1 appears to serve a gate keeper function, maintaining cellular homeostasis by restricting essential cytoplasmic processes to the cytoplasm(6). Crm1 cargos include a long list of tumor suppressors and oncogenic proteins, many of which are mislocalized in cancer cell types(109, 110). A firmer structural understanding of cargo recognition features such as provided by the HIV NEC structure may be expected to facilitate the development of specific Crm1 inhibitors.

## ACKNOWLEDGEMENTS

We thank all members of Frankel and Cheng labs for helpful comments and discussion. This work was supported by NIH/NIAID funding for the HIV Accessory and Regulatory Complexes (HARC) Center (U54AI170792 to A.D.F. and Y.C.). Instruments at the UCSF Cryo-EM facility are partially supported by grants from the NIH/OD (S10OD020054, S10OD021741 and S10OD026881) and Howard Hughes Medical Institute. Y.C. is an investigator of the Howard Hughes Medical Institute. David Bulkley, Glenn Gilbert, and Eric Tse from the UCSF EM core facility for technical assistance. Bhargavi Jayaraman and David Booth for bacterial and mammalian expression vectors.

## Methods

### Expression and protein purification

#### Expression and Purification of wild type human Crm1

Full-length and codon optimized human Crm1 was cloned into a modified pFastBac vector containing a mammalian cytomegalovirus (CMV) promoter between the BamHI and NotI restriction sites(111–113). Crm1 has an N-terminal octa-histidine tag followed by the maltose binding protein (MBP) separated by a Tobacco Etch Virus (TEV) protease site. This plasmid was transformed into the DH10Bac *Escherichia coli* strain to generate the recombinant bacmid DNA which was then transfected into insect cells to generate the BacMam virus(114). Crm1 was expressed in FreeStyle HEK293-F cells suspended in FreeStyle 293 Expression Medium (Invitrogen) supplemented with 2% (v/v) fetal bovine serum (FBS). Cells were grown at 37°C and 8% CO_2_ on an orbital shaker to a cell density of ∼2.0 x 10_6_ per mL at which point they were transduced with P3 BacMam virus and expression was enhanced by the addition of sodium butyrate to a final concentration of 10 mM. Cells were harvested 30 hours post-transduction and snap-frozen in liquid nitrogen and stored at -80°C until purification. Crm1 was resuspended in 5 mL of lysis buffer (20 mM Tris pH 7.5, 600 mM NaCl_2_, 2 mM magnesium acetate (MgOAc), 2 mM β-mercaptoethanol (BME), 10% glycerol (v/v) and 1 mM guanosine diphosphate (GDP)) for each gram of cell pellet. Lysis buffer was supplemented with Halt_TM_ Protease Inhibitor (Thermo Scientific) and phenylmethylsulfonyl fluoride (PMSF) to a final concentration of 1 mM. Crm1 was purified by batch binding to amylose resin (New England Biosciences, US) overnight at 4°C. The amylose resin was washed in Buffer A (20 mM Tris pH 7.5, 600 mM NaCl_2_, 2 mM MgOAc, 2 mM BME, 20% glycerol (v/v) and 20 mM Imidazole pH 7.5) followed by elution in Buffer A supplemented with 10 mM Maltose. The eluted Crm1 was TEV cleaved at room temperature. The TEV protease and the His_8_ – MBP tag were purified away from Crm1 using Ni-NTA resin. Crm1 was then exchanged into Buffer B (20 mM Tris pH 7.5, 100 mM NaCl_2_, 2 mM MgOAc, 2 mM BME and 20% glycerol (v/v)) and loaded onto a Mono-Q 5/50 GL column (Cytivia, Sweden). Crm1 was eluted with a gradient of 0-100% Buffer C (20 mM Tris pH 7.5, 1 M NaCl_2_, 2 mM MgOAc, 2 mM BME, 20% glycerol (v/v)). Fractions were analyzed by SDS-PAGE, pure fractions were pooled, frozen in liquid nitrogen and stored at -80°C. Crm1 concentration was determined by Bradford Assay with a bovine serium albumin (BSA) standard (Thermo Scientific).

#### Expression and Purification of Binding Site Variants of human Crm1

Full-length human Crm1 with an N-terminal 2x Strep tag was cloned into the pcDNA 4/TO plasmid between the restriction sites NotI and XhoI. The Crm1 variants were generated by site directed mutagenesis. The Crm1 variants were expressed using the Expi293 transient expression system (ThermoFisher, US) using manufacturer supplied protocols. Cells were harvested 48 hours post-transfection and snap-frozen in liquid nitrogen and stored at -80°C until purification. Crm1 was resuspended in 5 mL of lysis buffer (20 mM Tris pH 7.5, 600 mM NaCl_2_, 2 mM MgOAc, 2 mM BME, 10% glycerol (v/v) and 1 mM GDP) for each gram of cell pellet. Lysis buffer was supplemented with Halt^TM^ Protease Inhibitor (Thermo Scientific) and PMSF to a final concentration of 1 mM. Crm1 was purified by batch binding to Strep-Tactin XT 4Flow resin (IBA, Germany) overnight at 4°C. The StrepTactin resin was washed in Buffer D (20 mM Tris pH 7.5, 600 mM NaCl_2_, 2 mM MgOAc, 2 mM BME and 20% glycerol (v/v)) followed by elution in Buffer D supplemented with 1x BXT Buffer (IBA, Germany). The eluted Crm1 was then exchanged into Buffer B and loaded onto a Mono-Q 5/50 GL column (Cytivia, Sweden). Crm1 was eluted with a gradient of 0-100% Buffer C. Fractions were analyzed by SDSPAGE, pure fractions were pooled, frozen in liquid nitrogen and stored at -80°C. Crm1 concentration was determined by Bradford Assay with a BSA standard (Thermo Scientific).

#### Expression and Purification of Rev Variants

Plasmids used for wild type Rev HIV-1 HXB3 and Rev L60R HIV-1 strain HXB3 have been previously described(40). Rev L81A HIV-1 strain HXB3 was generated by site directed mutagenesis. Rev NS2Val was generated by modifying the previously described Rev construct through PCR sewing with primers that amplified Rev on either side of the native NES adding 13 amino acids to both the N-terminal and C-terminal halves. The two PCR products contained a 38 bp overlap aiding in the PCR amplification of the full Rev NS2Val construct that could then be cloned into the original plasmid through restriction site digestion (using the NdeI and XhoI sites) followed by ligation to replace wild-type Rev with the Rev tight NES variation. This increased the rev gene from 116 amino acids to 121 amino acids. The above Rev variants were expressed as previously described(40). Cells were harvested and snap-frozen in liquid nitrogen and stored at -80°C until purification. Genes blocks of the Rev linked dimer variants were designed to have the same Rev DNA sequence from the Rev wild type plasmid used in our study in the N-terminal position followed by a (GGGGS)3 linker and a codon optimized DNA sequence for Rev HIV-1 strain HXB3 in the C-terminal position. For the 11A/11GS construct residues 62-72 were replaced with 11 alanines in the N-terminal Rev and for the C-terminal Rev theses 11 residues were replaced with (GGS)3GS. The gene fragments (IDT) were designed to have a 38 bp overlap at the 5’ and a 16 bp overlap at the 3’. The wildtype plasmid was digested using the enzymes NdeI and XhoI and gel purified. The fragments and the digested plasmid were then combined using In-Phusion Snap Assembly (Takara Bio USA). The linked dimer constructs were expressed in *E. coli* strain BL21/DE3. Cells were grown to OD_600_ = 0.6-0.8 at 37°C in 2XYT media (Fisher Scientific, US) with 100 μg/mL carbenicillin. Cells were then cooled to 18°C, Isopropyl-β-D-thiogalactopyranoside (IPTG) was added to 1 mM to induce expression. Cells were harvested the following day and snap-frozen in liquid nitrogen and stored at -80°C until purification. To purify each Rev protein was resuspended in 5 mL of lysis buffer (50 mM Tris pH 7.5, 250 mM NaCl_2_, 5 mM BME, 10 mM imidazole pH 7.5, 0.1% (v/v) Tween-20) per gram of cell pellet. Lysis buffer was supplemented with Halt^TM^ Protease Inhibitor (Thermo Scientific), PMSF to a final concentration of 1 mM and lysozyme to a final concentration of 0.5 mg/mL. Endogenous RNA from *E. coli* was removed by the addition of solid NaCl_2_ (final concentration of 2 M) and urea (final concentration of 1M) to the cleared lysate before addition to Ni-NTA resin. Batch binding was carried out at 4°C for 1 hour. The Ni-NTA resin was washed with Buffer E (50 mM Tris pH 7.5, 2 M NaCl_2_, 5 mM BME, 10 mM imidazole pH 7.5, 0.1% (v/v) Tween-20, 1M Urea) and Buffer F (125 mM Phosphate buffer pH 7.5, 250 mM NaCl_2_, 5 mM BME, 10 mM imidazole). Rev was eluted by stepwise chromatography in Buffer F (50-500 mM imidazole pH 7.5). Fractions were analyzed by SDS-PAGE, pure fractions were pooled, frozen in liquid nitrogen and stored at -80°C. Rev concentration was determined by Bradford Assay with a cysteine free construct of Rev as a standard. Prior to addition with the RRE, Rev was dialyzed overnight in Buffer F (10 mM imidazole pH 7.5) supplemented with 10% (v/v) glycerol in the presence of TEV protease. The N-terminal hexa-histidine tag and GB1 solubility-enhancing domain and the N-terminal hexa-histidine tagged TEV protease was purified away from the TEV cleaved Rev by Ni-NTA resin.

#### Ran Q69L 180 Expression and Purification

Ran Q69L 180 was generated by site directed mutagenesis to add a stop codon in place of alanine 181 to the previously described Ran plasmid(23). Ran Q69L 180 was expressed as previously described(23). Cells were harvested and snap-frozen in liquid nitrogen and stored at -80°C until purification. Ran Q69L 180 was resuspended in 5 mL of lysis buffer (64 mM Tris pH 7.5, 400 mM NaCl_2_, 5 mM MgCl2, 2 mM BME, 10% (v/v) glycerol) per gram of cell pellet. Lysis buffer was supplemented with Halt^TM^ Protease Inhibitor (Thermo Scientific), PMSF to a final concentration of 1 mM and lysozyme to a final concentration of 0.5 mg/mL. RanQ69L 180 was first purified by stepwise elution from Ni-NTA in Buffer G (64 mM Tris pH 7.5, 400 mM NaCl2, 5 mM MgCl2, 2 mM BME, 10% (v/v) glycerol, 50-500 mM imidazole pH 7.5). Following Ni-NTA purification Ran Q69L 180 was dialyzed overnight in Buffer G (10 mM imidazole pH 7.5) at 4°C in the presence of N-terminal hexa-histidine tagged TEV protease. The TEV protease and the deca-histidine tag were purified away from the TEV cleaved Ran Q69L 180 by Ni-NTA resin. Prior to cation exchange Ran Q69L 180 was concentrated with a 10 kD MWCO Amicon Centrifugal Filter at 3.5K x g to a final volume of 2.5 mL. Concentrated Ran Q69L 180 was exchanged into Low Salt Buffer H (25 mM HEPES pH 7.5, 100 mM KCl, 4 mM BME, 10% (v/v) glycerol, 30 μM GTP). Cation exchange was performed on a Mono-S column 5/50 GL column (Cytivia, Sweden) with a linear gradient 0-100% High Salt Buffer I (25 mM HEPES pH 7.5, 1 M KCl, 4 mM BME, 10% (v/v) glycerol, 30 μM GTP). Fractions were analyzed by SDS-PAGE, pure fractions were pooled, frozen in liquid nitrogen and stored at -80°C. RanQ69L 180 concentration was determined by Bradford Assay with a BSA standard (Thermo Scientific).

#### _MBP_Nup_1916-2033_ FG Peptide Expression and Purification

_MBP_Nup_1916-2033_ was cloned into pET22b between the NdeI and XhoI cut sites. This construct contains an N-terminal MBP tag followed by 118 residues of Nup214 (residues 1916-2033), the amino acids leucine and glutamate (LE) and a C-terminal hexa-histidine tag. _MBP_Nup214_1916-2033_ was expressed in Rosetta^TM^(DE3) pLysS cells. Cells were grown to OD_600_ = 0.6-0.8 at 37°C in 2XYT media (Fisher Scientific, US) with 100 μg/mL carbenicillin. Cells were then cooled to 16°C and expression was induced with IPTG to a final concentration of 0.5 mM. Cells were harvested after 20 hours and snap-frozen in liquid nitrogen and stored at -80°C until purification. One liter of _MBP_Nup214_1916-2033_ cell pellet was resuspended in 80 mL of lysis buffer (50 mM HEPES pH 7.4, 500 mM NaCl_2_, 5% (v/v) glycerol, 20 mM Imidazole and 1 mM 1,4-Dithiothreitol (DTT)). Cleared lysate was added to equilibrated Ni-NTA resin, sample was flowed over resin twice. The resin was then washed with a 100 mL of lysis buffer. _MBP_Nup214_1916-2033_ was eluted from the Ni-NTA resin with 20 mL of Buffer J (50 mM HEPES pH 7.4, 500 mM NaCl_2_, 5% (v/v) glycerol, 500 mM Imidazole and 1 mM DTT). _MBP_Nup214_1916-2033_ was then added to amylose resin. Batch binding was carried out at 4° rotating end over end for 1 hour. Resin was then washed with 60 mL of Buffer K (20 mM Tris pH 7.4, 200 mM NaCl_2_, 1 mM ethylenediaminetetraacetic acid (EDTA), 2 mM DTT). Protein was eluted with 20 mL of Buffer K supplemented with Maltose to a final concentration of 10 mM. Fractions were analyzed by SDS-PAGE, pure fractions were pooled, frozen in liquid nitrogen and stored at -80°C. _MBP_Nup214_1916-2033_ concentration was determined by Bradford Assay with a BSA standard (Thermo Scientific).

#### Expression and Purification of Human Snurportin

Full-length human Snuportin (SPN) with an N-terminal 2x Strep tag was cloned into the pcDNA 4/TO plasmid between the restriction sites NotI and XhoI. SPN was expressed using the Expi293 transient expression system (ThermoFisher, US) using manufacturer supplied protocols. Cells were harvested 48 hours post-transfection and snap-frozen in liquid nitrogen and stored at -80°C until purification. SPN was resuspended in 5 mL of lysis buffer (50 mM Tris pH 7.5, 200 mM NaCl_2_, 2 mM DTT, 2 mM MgOAc and 1 mM EDTA) for each gram of cell pellet. Lysis buffer was supplemented with Halt^TM^ Protease Inhibitor (Thermo Scientific) and PMSF to a final concentration of 1 mM. SPN was purified by batch binding to Strep-Tactin XT 4Flow resin (IBA, Germany) overnight at 4°C. The Strep-Tactin resin was washed in lysis buffer followed by elution in lysis buffer supplemented with 1x BXT Buffer (IBA, Germany). The eluted SPN was then exchanged into Buffer L (50 mM Tris pH 7.5, 50 mM NaCl2, 2 mM DTT and 2 mM MgOAc) and loaded onto a Mono-Q 5/50 GL column (Cytivia, Sweden). SPN was eluted with a gradient of 0-100% Buffer M (50 mM Tris pH 7.5, 1 M NaCl_2_, 2 mM DTT and 2 mM MgOAc). Fractions were analyzed by SDS-PAGE, pure fractions were pooled, frozen in liquid nitrogen and stored at -80°C. SPN concentration was determined by Bradford Assay with a BSA standard (Thermo Scientific).

#### RNA Transcription and Purification

The plasmid used for RRE234 HIV-1 strain SF-2 has been previously described(23). The plasmid for RRE61 was generated by site directed mutagenesis on the RRE234 plasmid making the G262A and G269A mutations (Figure S10). A gene block was designed for RRE355 HIV-1 strain SF-2 with the restriction sites SacII and EcoRI at the 5 prime end and XbaI at the 3 prime end. RRE234 SF-2 was then digested out of the previously described plasmid and RRE355 SF-2 was ligated in. Plasmids were linearized utilizing the EcoRI site. RNA was transcribed by T7 polymerase from the linearized plasmid followed by purification by denaturing Urea-PAGE(23, 40). The pure RNA pellets were resuspended in 0.5 mM HEPES-KOH pH 7.5. RNA was annealed by heating to 95-100°C then flash cooled on ice. RNA was quantified by UV spectroscopy using a 260 nm extinction coefficient of 2.5 x 10^6^ for RRE234 and RRE61 and a 260 nm extinction coefficient of 4.3 x 10^6^ for RRE355.

#### Nuclear Export Complex Assembly

Rev and RRE were mixed in an 8:1 ratio for the monomer constructs and 4:1 ratio for the linked dimer constructs. The RNP mixture was then loaded onto a Superdex 200 Increase 10/300 (Cytiva, Sweden). The RNP complex was purified in Buffer N (50 mM HEPES pH 7.5, 50 mM KCl, 2 mM BME, 2% (v/v) glycerol). The RNP was then mixed with RanQ69L 180, Crm1 and _MBP_Nup214_1916-2033_ in a ratio of 1:8:4:8 in Buffer O (50 mM HEPES pH 7.5, 100 mM KCl, 2 mM MgOAc, 2 mM BME, 30 μM GTP). The complex was then incubated at 37°C for 15 minutes. Post incubation the complex was kept at room temperature until it was added to the grid. Complexes assembled for negative stain only did not contain _MBP_Nup214_1916-2033_. The complex was quantified by UV spectroscopy using a 260 nm extinction coefficient of 2.5 x 10^6^ for RRE 234 and RRE 61 and a 260 nm extinction coefficient of 4.3 x 10^6^ for RRE355.

#### Negative-stain Electron Microscopy

EM grids of negatively stained hCrm1 alone, NEC or hCrm1/SPN were prepared by following the established protocol(115). Briefly, 3 uL of sample at ∼0.010–0.012 mg/mL was applied to a glow-discharged copper grid layered with amorphous carbon (Ted Pella Inc., US). After 30 seconds the sample was blotted off with filter paper (Whitman, #1) and washed twice using 3 μL of 2% (w/v) uranyl formate that was blotted off with filter paper after each application. Negatively stained grids were imaged on either a FEI Tecnai T12 electron microscope (Thermo Fisher Scientific, US) equipped with a LaB6 filament and a 4K CCD camera (UltraScan 895, Gatan, Inc.), operated at 120 kV, a FEI Tecnai T20 electron microscope (Thermo Fisher Scientific, US) equipped with a LaB6 filament and a TVIPS TemCam F816 (8K x 8K) scintillator-based CMOS camera (TVIPS, Germany), operated at 200 kV or a Talos L120C (Thermo Fisher Scientific, US) equipped with a LaB6 filament and a Ceta-D camera (Thermo Fisher Scientific, US) operated at 120 kV. Particle picking and 2D classification was carried out using RELION(116, 117).

#### Cryo-EM Grid Preparation

Functionalized graphene oxide (GO) grids(118, 119) were used to mitigate complex disassociation and preferred orientation as observed with non-modified grids or GO alone treated grids. GO grids were functionalized following the established protocol(120) with a variety of single-stranded DNA (ssDNA) oligos of various lengths and sequences or polyamine (Table S3)(121, 122). Briefly, first GO was applied to 300 mesh 1.2/1.3 R Au Quantifoil grids (Qunatifoil, Germany) that were plasma cleaned in argon gas for 10 seconds using an established protocol(119). To functionalize GO grids with polyamine, 5 μL of ethylenediamine was added to 5 mL of dimethyl sulfoxide (DMSO). The GO grids were incubated in this solution for ∼3 hours. The ethylenediamine/DMSO mixture was aspirated off and the grids were then incubated for 1 minute in 5 mL of DMSO followed by aspiration. Next the grids were incubated for 1 minute in deionized water (ddH2O), three times before being incubated in ethanol for 1 minute three times. After this they were allowed to air dry before either being used or stored at -20°C. For ssDNA functionalization, DNA oligos were first resuspended in DMSO to a final concentration of 200 μM. Single grids were floated GO side down on 20 μL of ss-DNA/DMSO solution in a 1.5 mL Eppendorf tube. The tubes were then placed in an Eppendorf thermomixer (Eppendorf, US) at 25°C overnight. The following day the grids were rinsed with 200 μL of ddH2O followed by 200 μL of ethanol. Grids were air dried before either being used or stored at - 20°C. Three μL of sample at ∼0.5-0.7 mg/mL was added to the functionalized grid. The sample was allowed to incubate on the grid for 1 minute before being side blotted off outside of the vitrobot, two applications of 3 μL of Buffer J were used to wash the grid after sample addition. After the addition of the last buffer drop the grid and the tweezers were placed in the vitrobot for immediate blotting and plunging. Grids were frozen using a FEI Vitrobot Mark IV (Thermo Fisher Scientific, US) with a blotting time of 1 second. Vitrobot was set to 100% humidity and 22°C. Grids were plunge-frozen in liquid ethane and cooled by liquid nitrogen.

#### Cryo-EM Data Acquisition

Data sets were acquired on a Titan Krios (Thermo Fisher Scientific, US), operated at 300 kV and equipped with a K3 direct electron detector and a BioQuantum energy filter (Gatan, US) with a slit set to 20 eV. A defocus range of 0.8 to 2 μm under focus was used. Super-resolution movies were collected semiautomatically using SerialEM and recorded with a super-resolution pixel size of 0.417 Å/pixel for the ssDNA-GO grids and 0.4175 Å/pixel for the polyamine-GO grids(123). A nominal magnification of 105kx was used for all samples, a total dose of 66 e-/Å^2^ over 115 frames was used for the ssDNA samples and a total dose of 68 e-/Å^2^ over 118 frames was used for the polyamine-GO samples. Of the 22,762 movies that were recorded for the polyamine-GO grids, 7,720 movies were collected at 0° tilt, 11,190 movies were collected at 15° tilt and 3,852 movies were collected at 30° tilt. Of the 24,447 movies that were recorded for the ssDNA-GO grids, 7,286 movies were collected at 25° tilt. Movies were motion corrected using MotionCor2 and binned 2 x 2 by Fourier cropping to a final pixel size of 0.834 Å per pixel for ssDNA samples and 0.835 Å per pixel for polyamine grids(124).

#### Image Processing

Initial processing for the ssDNA samples was performed in CryoSPARC(125). Dose-weighted motion corrected sums for the ssDNA samples were used for the estimation of the contrast transfer function (CTF) utilizing Patch CTF in CryoSPARC. Particles were auto picked individually for all six of the ssDNA samples using a Gaussian template in CryoSPARC. Particles were extracted with a box size of 440 pixels and binned to 220 pixels. 2D classification was carried out in CryoSPARC. Particles from the different collections were combined and 2D class averages were used to re-picked particles by the template-based method in CryoSPARC. 2D classification was ran multiple times to remove particle images that did not resemble the NEC. Particle files were converted to RELION using UCSF pyEM(126) for subsequent image processing. Image processing for the polyamine samples was performed in RELION(117). Dose-weighted motion corrected sums were used for the estimation of the CTF utilizing CTFFIND4 wrapped within the RELION graphical user interface(127). Particles were picked using a 3D template from the combined results from the ssDNA samples. Particles were extracted with a box size of 440 pixels and binned to 110 for 2D classification. After multiple rounds of 2D classification, selected particles were reextracted in RELION and binned to 220 pixels. Particles from 2D classification from the different functionalized grids and tilt angles were combined and 3D classification and refinement were carried out in RELION. Employing symmetry expansion and 3D variability methods(128) led to a subset of particles with a sufficient sampling of angular space (Figure 2E)(116, 117, 125). A flow chart of image processing is illustrated in Figure S11. Local filtering of the final map was carried out in CryoSPARC, map sharpening was carried out using DeepEMhancer(129). Resolution was determined from Fourier Shell Correction (FSC) using criterion of FSC = 0.143(130). Map to model FSC curves were determined with the program EMDA(131), with a resolution criterion of FSC = 0.5 In addition, directional FSC (dFSC)(132) was used to evaluate the directional uniformity of the reconstructions, as reported in Table S1. Figures were generated from the map that is displayed with a spatially adaptive filter generated with a local resolution map (CryoSPARC)(128).

#### Model Building and Refinement

The atomic models of human Crm1/Ran-GTP (PDB: 5DIS)(53) and Rev/RRE (PDB: 4PMI)(54) were docked into the cryo-EM map as a rigid body using UCSF ChimeraX(133). Residues 389-400 of Crm1 and residues 61-91 of Rev were modeled using Rosetta Enumerative Build(134) followed by real space refinement(55). Except for Crm1 residues 389-400, Crm1 and Ran are undistinguishable from the previously solved crystal structure(53). We utilized the Rosetta program DRRAFTER(135) to model the 39 nucleotides (119-154, 171-174) of the RRE that are ordered in the NEC, inputting the stem IIB portion of the RRE from the Rev/RRE dimer co-crystal structure(54) together with RNA SHAPE data(42). The first 11 and the last 30 amino acids of Rev were not observed. The _MBP_Nup214_1916-2033_ peptide is observable at higher contour levels and rigid body docking of the crystal structure of Crm1-SPN-RanGTP-_MBP_Nup214 that the Nup214 peptide is bound to each Crm1 subunit (S12)(53). The modeled complex went through multiple rounds of refinement in real space using Phenix and manual adjusting using COOT(55, 136). Map-to-model FSC plot was generated using EMDA(131). Visualizations of the atomic model were generated in UCSF ChimeraX(133).

#### Dual Luciferase Assays

Rev reporter activity was determined by a fluorescence-based assay utilizing HEK293T cells transfected with Revand RRE234-containing reporter constructs. HEK293T cells in a 96-well plate were co-transfected with 0.5 ng of Rev wild-type or mutant plasmid, 49.5 mg of RRE234 reporter plasmid (pCMV gagpol-IRES FFL-RRE) and 0.5 ng of the reference plasmid pNL1.1.TK [Nluc/TK] (Promega, N1501) with PolyJet (SignaGen) following the manufacturer’s instructions. Each sample was performed in biological triplicate. After 24 hours of transfection, the cells were lysed in 1x Passive Lysis Buffer (Promega), firefly luciferase and NanoLuc luciferase activities were measured with the Promega Nano-Glo® Dual-Luciferase® Reporter Assay System and microplate reader (Ultra Evolution, TECAN). Rev-dependent firefly luciferase values were normalized to NanoLuc values. Rev expression levels were verified by Western blot using Flag antibody (Sigma, F1804) with α-Tubulin (Sigma, T5168) used as a loading control. IRES-FFL gene was inserted into pCMV gagpol-RRE234 vector as previously described by Gibson Assembly for testing in Rev reporter assays(77). Rev-Crm1 reporter activity was determined by a fluorescence-based assay utilizing BHK21 cells transfected with Crm1 wild-type or mutations and Rev-RRE containing reporter constructs. BHK21 in a 96-well plate were transfected with 20 ng Crm1 wild-type or mutant, 4 ng of Rev, 99 ng of RRE reporter plasmid (pCMV gagpol-IRES FFL-RRE) and 1 ng of the reference plasmid pNL1.1.TK [Nluc/TK] (Promega, N1501) were co-transfected with Lipo-fectamine™ 3000 (Invitrogen™) following the manufacturer’s instructions. Each sample was performed in biological quadruplicates. After 24 hours of transfection, the cells were lysed in 1x Passive Lysis Buffer (Promega), firefly luciferease and NanoLuc luciferase activities were measured with the Promega Nano-Glo® Dual-Luciferase® Reporter Assay System and microplate reader (Ultra Evolution, TECAN). Rev-dependent firefly luciferease values were normalized to NanoLuc values. Proteins were detected with indicated antibodies.

#### Co-Immunoprecipitation Assays

Expi293 cells were lysed in lysis buffer (50 mM Tris pH 7.5, 150 mM NaCl_2_, 2 mM MgCl_2_, 0.5% Triton X-100 (v/v) 0.2% Na-deoxycholate (w/v)) supplemented with HaltTM Protease Inhibitor (Thermo Scientific) for 30 minutes at 4°C by gentle rocking. Cleared cell lysates were incubated over night at 4°C with Strep-Tactin XT 4Flow resin (IBA, Germany). The Strep-Tactin XT 4Flow resin was washed with lysis buffer without Protease Inhibitor. Proteins were eluted with Buffer P (50 mM Tris pH 7.5, 150 mM NaCl_2_, 2 mM MgCl_2_, 0.5% Triton X-100 (v/v), 1x BXT Buffer (IBA, Germany)). Eluate was analyzed by Western blotting with antibodies against the Flag-tag, 2x Strep-tag or Crm1.

#### Gel Shift Assays

Electrophoretic mobility shift assays were performed in Buffer Q (10 mM HEPES-KOH pH 7.5, 100 mM KCl, 1 mM MgCl_2_, 0.5 mM EDTA, 10% glycerol (v/v), 0.2mg/mL BSA, 50 ug/mL yeast tRNA). Crm1 protein stock for RNA binding was prepared in Buffer Q supplemented with twenty-five-fold molar excess of yeast tRNA at twice the final concentration (1.6 uM). Two-fold serial dilutions of Crm1 were made for six reactions (0-1.6 uM). RRE RNA was diluted in Buffer Q at twice the final concentration (200 nM). Equal volumes of Crm1 and RRE RNA were mixed, and reactions were incubated at room temperature for 15 min before being loaded onto continuously running 8% nondenaturing polyacrylamide gels (0.5x Tris-Borate-EDTA). Gels were run at room temperature for 1-2 hours, stained with SYBR gold and visualized under UV light.

## Author Contributions

A.D.F. and Y.C. conceived the project and supervised the research. A.M.S. designed and expressed purified protein complexes, prepared EM grids (including GO application and functionalization), collected and processed all EM data and built and refined the model. A.M.S., A.V. and Y.L. cloned the mammalian plasmids used for the dual luciferase assays. Y.L. conducted all the dual luciferase assays and western blots. A.M.S. and A.V. cloned the bacterial plasmids. All authors participated in discussion and analysis of the data. A.D.F., Y.C. and A.M.S. prepared the manuscript.

## Data and Code Availability

All data generated or analyzed during this study are included in this published article and its Supplementary Information. The cryo-EM density map has been deposited in the Electron Microscopy Data Bank under accession code EMD-42494 and the coordinates have been deposited in the Protein Data Bank under accession 8URJ.

## Supplementary Information

**Figure S1.**
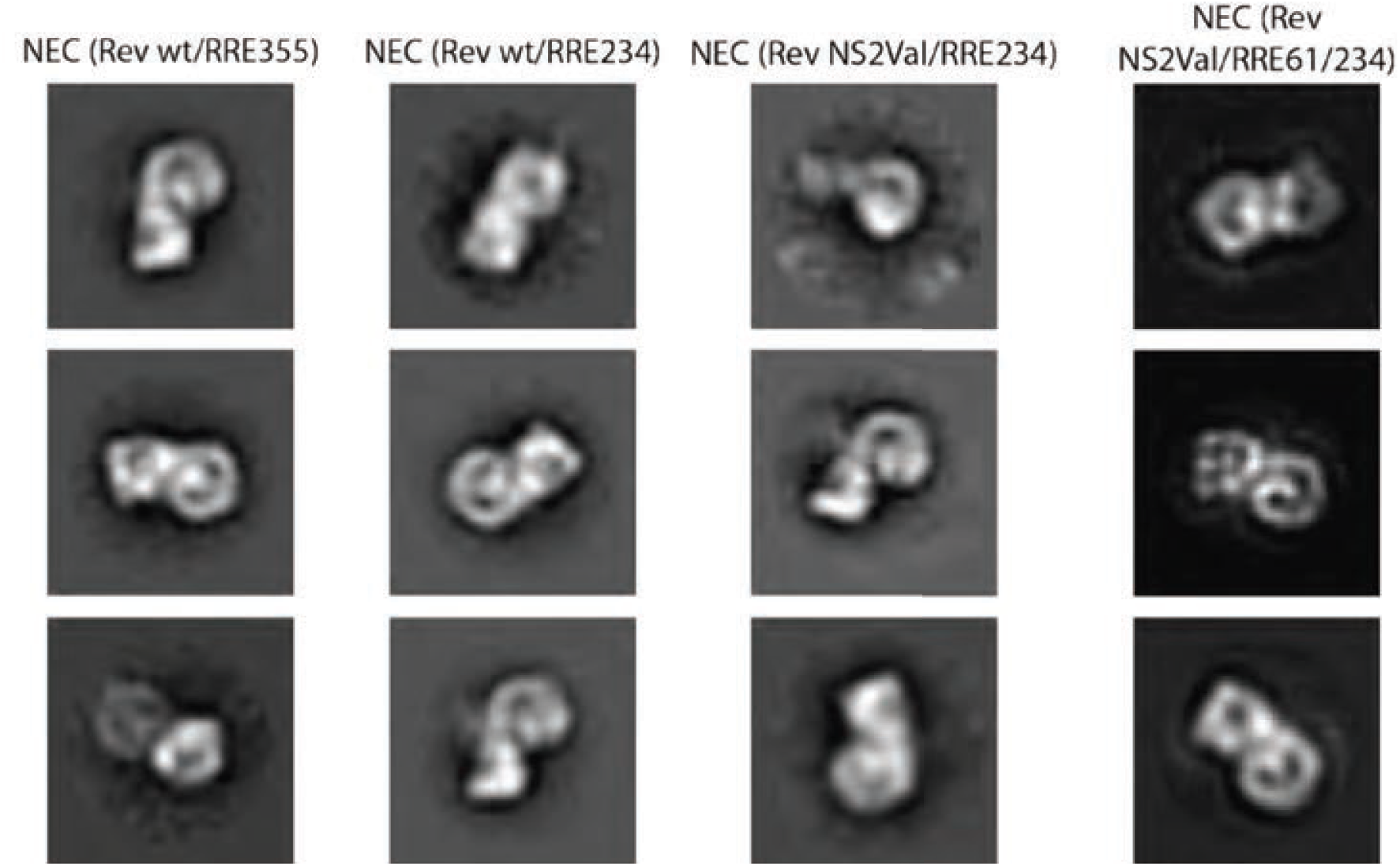
Representative negative stain 2D class averages of nuclear export complexes assembled with the different Rev and RRE variants used in our study. Complexes made with either wild-type Rev or Rev NS2Val and either RRE355, RRE234 or RRE61.

**Figure S2.**
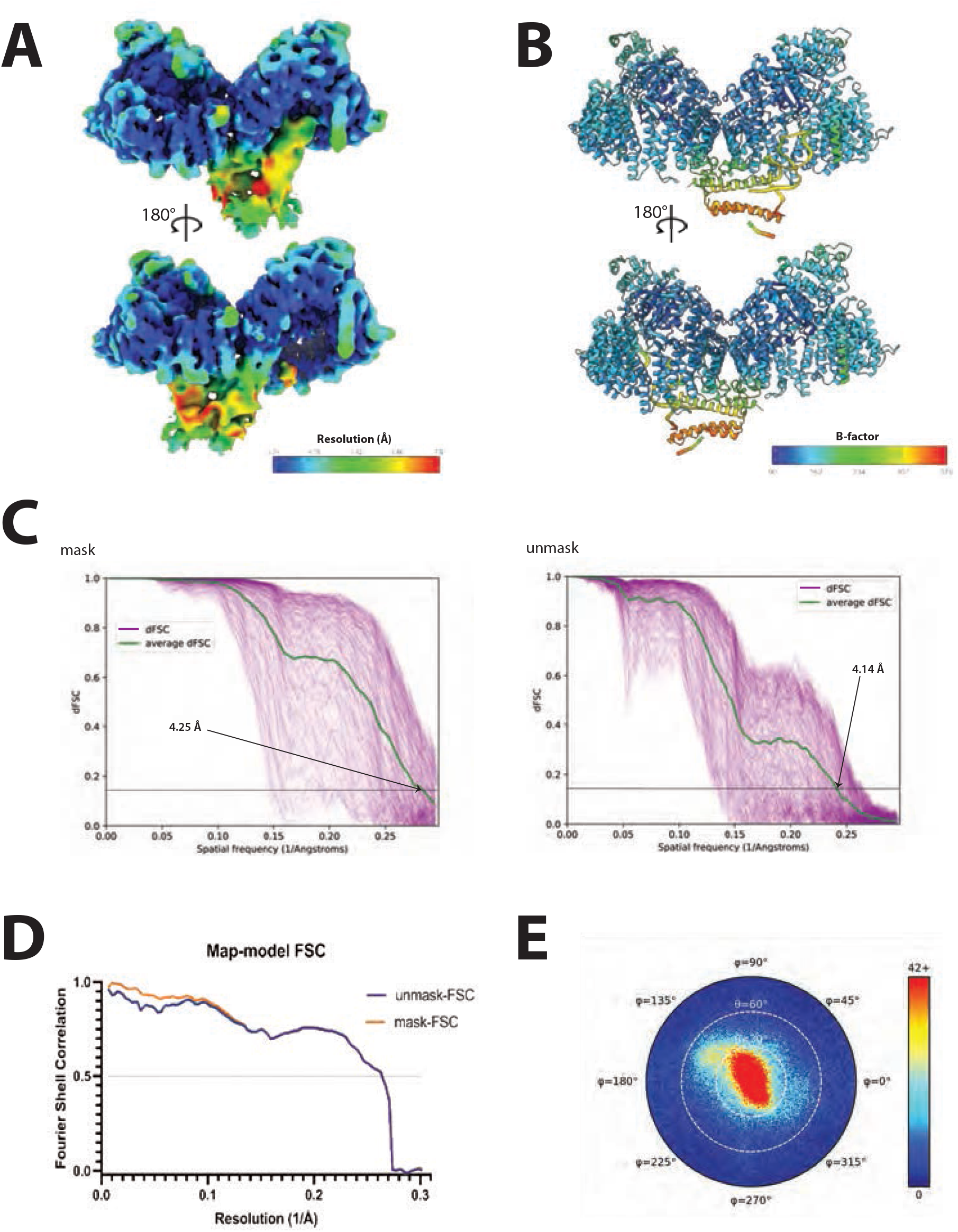
Cryo-EM map quality and metrics. (A) Local resolution estimate of the HIV-1 NEC reconstruction (cryoSPARC, local filtered map). (B) Refined NEC model colored according to *B*-factor. (C) Directional FSC curves were estimated as previously described^121^. Left is with a mask, right is without. (D) Map to model FSC calculated by EMDA. (E) Euler angle distribution of all particles used in the final 3D reconstruction (cisTEM^127^).

**Figure S3.**
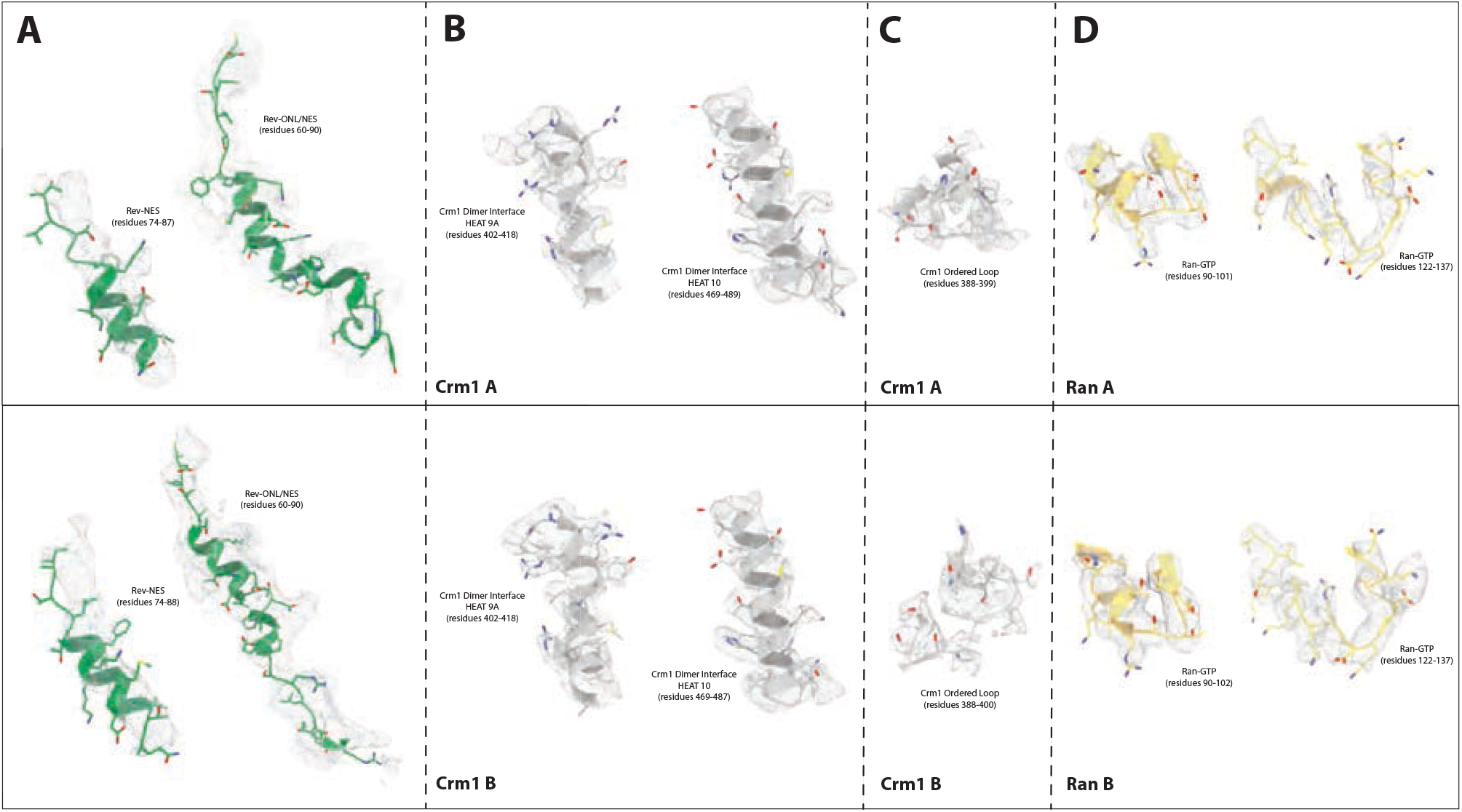
Representative cryo-EM densities from selected structural features. Representative segments of our cryo-EM map for various structural features highlighted in the main text. (A) Top: Rev NES and NES/ONL that interacts with Crm1 A. Bottom: Rev NES and NES/ONL that interacts with Crm1 B. Map of NES alone is the sharpened map from DeepEMHancer. Map of ONL/NES is the locally filtered map from CryoSPARC. Panels B-D: Top: Crm1 A or Ran-GTP A. Bottom: Crm1 B or Ran-GTP B. (B) The two helices that make up the Crm1 dimer interface and contain the species-specific residues (sharpened map from DeepEMHancer). (C) The ordered loop of Crm1 (residues 388-400) that becomes ordered in the presence of the Rev/RRE RNP. Map is the locally filtered map from CryoSPARC. (D) Loops from Ran-GTP that contain the residues (R95, K132 and K134) that interact with stem IIB (sharpened map from DeepEMHancer).

**Figure S4.**
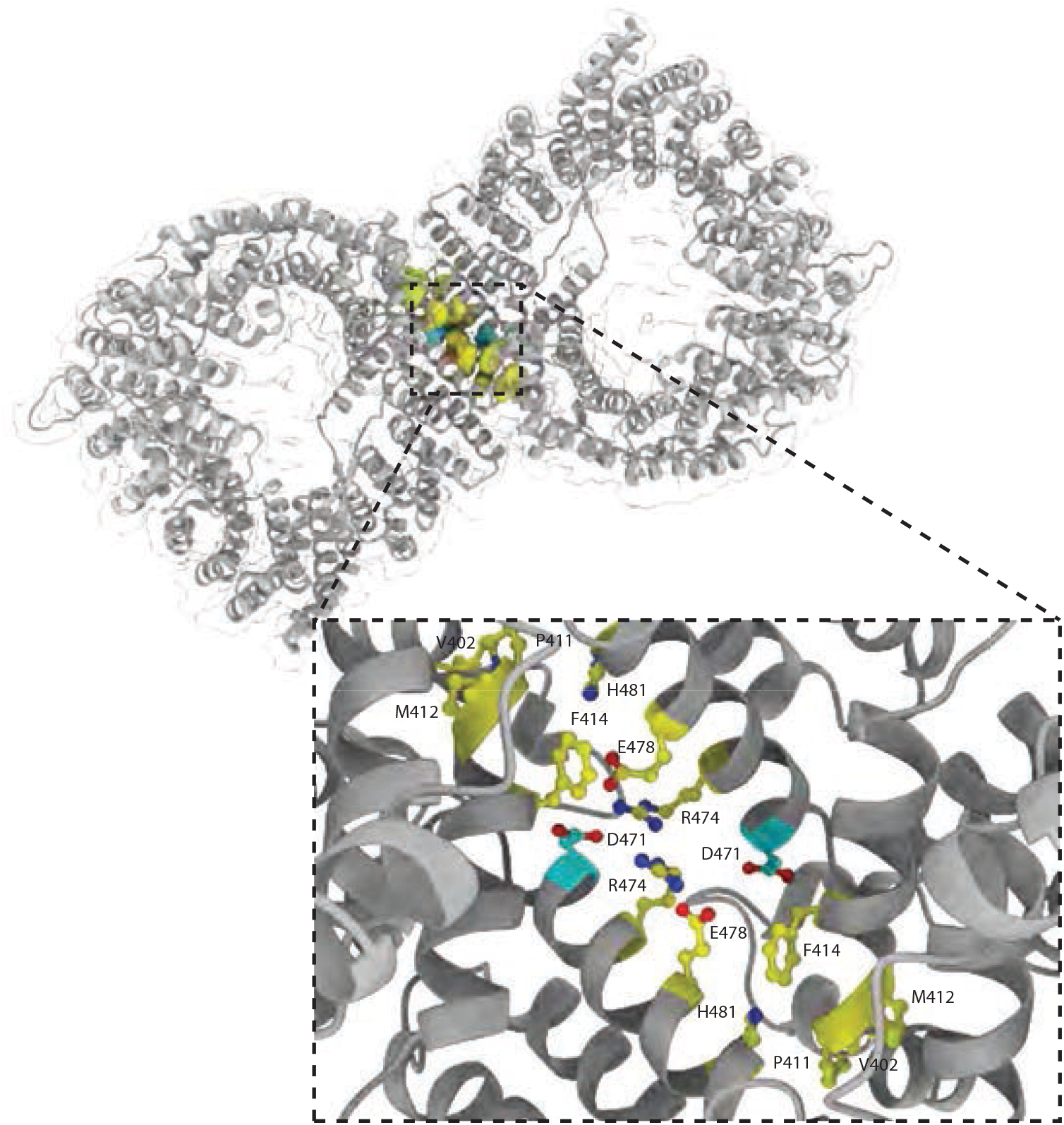
Crm1 dimer interface with species-specific residues. Top: Ribbon diagram of the hCrm1 dimer (gray) with the higher primate species-specific residues critical for HIV-1 replication shown in yellow and a conserved aspartate residue shown in cyan. Bottom: Close-up view of the hCrm1 dimer interface and the polar network created by the species-specific residues (V402, P411, M412, F414, R474, E478 and H481; shown in yellow) in combination with a conserved aspartate residue (D471; shown in cyan).

**Figure S5.**
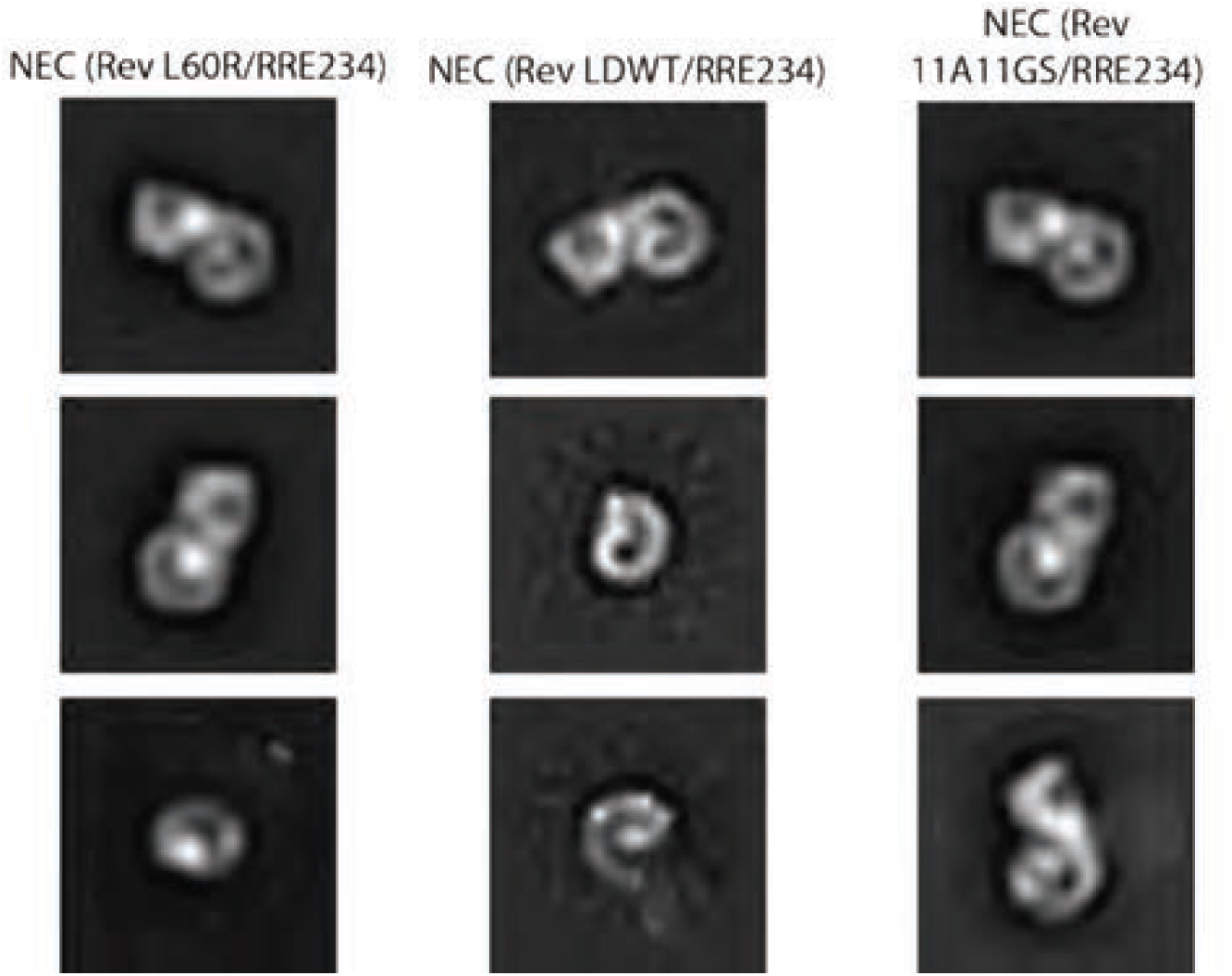
Representative negative stain 2D class averages of Rev variants to test efficiency of NEC assembly. NECs were assembled with either the monomeric oligomerization deficient mutant (L60R), the linked wild-type dimer (LDWT) or the linked Rev dimer with a helical ONL in the N-terminal position and a flexible ONL in the C-terminal position (11A/11GS). All NEC complexes were assembled with HIV-1 RRE234.

**Figure S6.**
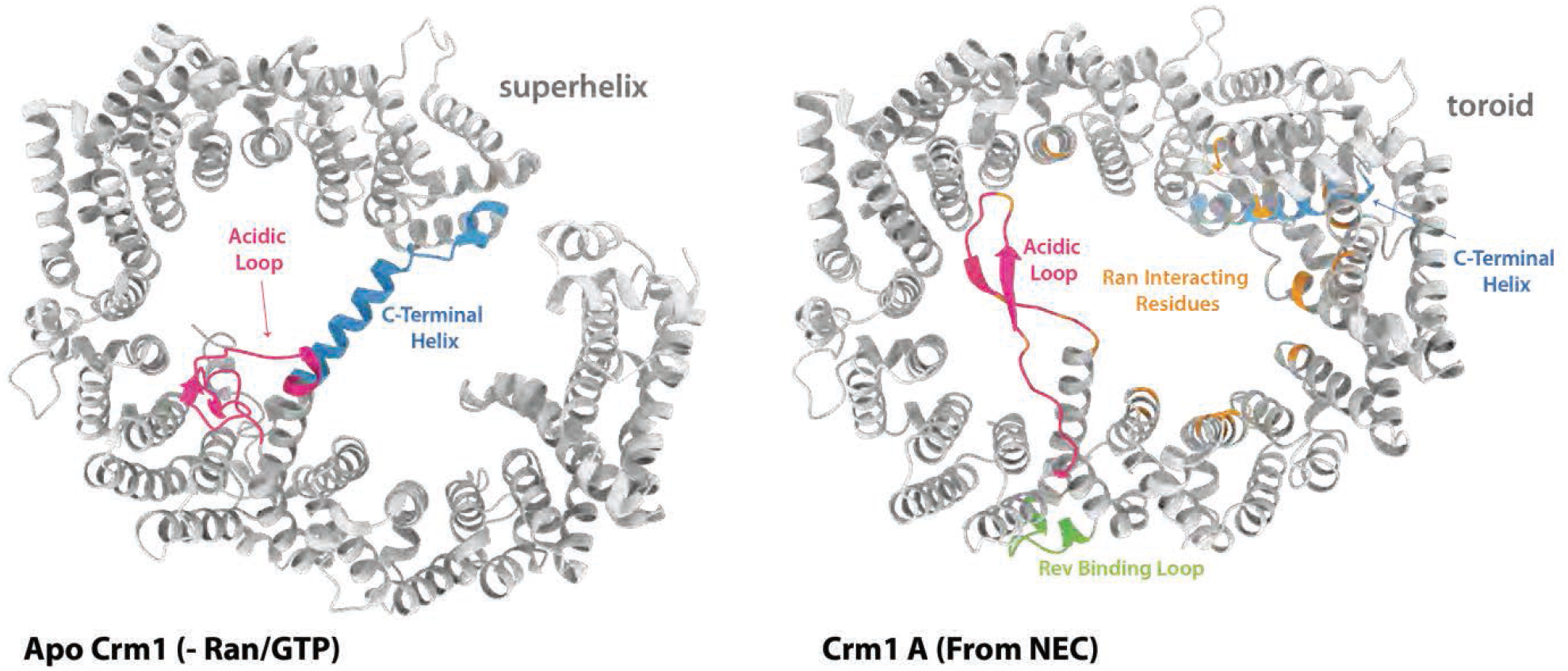
Structures of apo Crm1 (-RanGTP and cargo) vs Crm1 from the HIV-1 NEC. Left: Apo Crm1 (PDB: 4FGV) without Ran-GTP or cargo shown in ribbon diagram in gray. The acidic loop that interacts with Ran-GTP shown in pink. The inhibitory C-terminal helix shown in blue. Right: Crm1 A from NEC shown in ribbon diagram/gray, acidic loop colored in pink, C-terminal inhibitory helix shown in blue, residues that interact with Ran-GTP shown in orange and the Rev binding loop shown in green.

**Figure S7.**
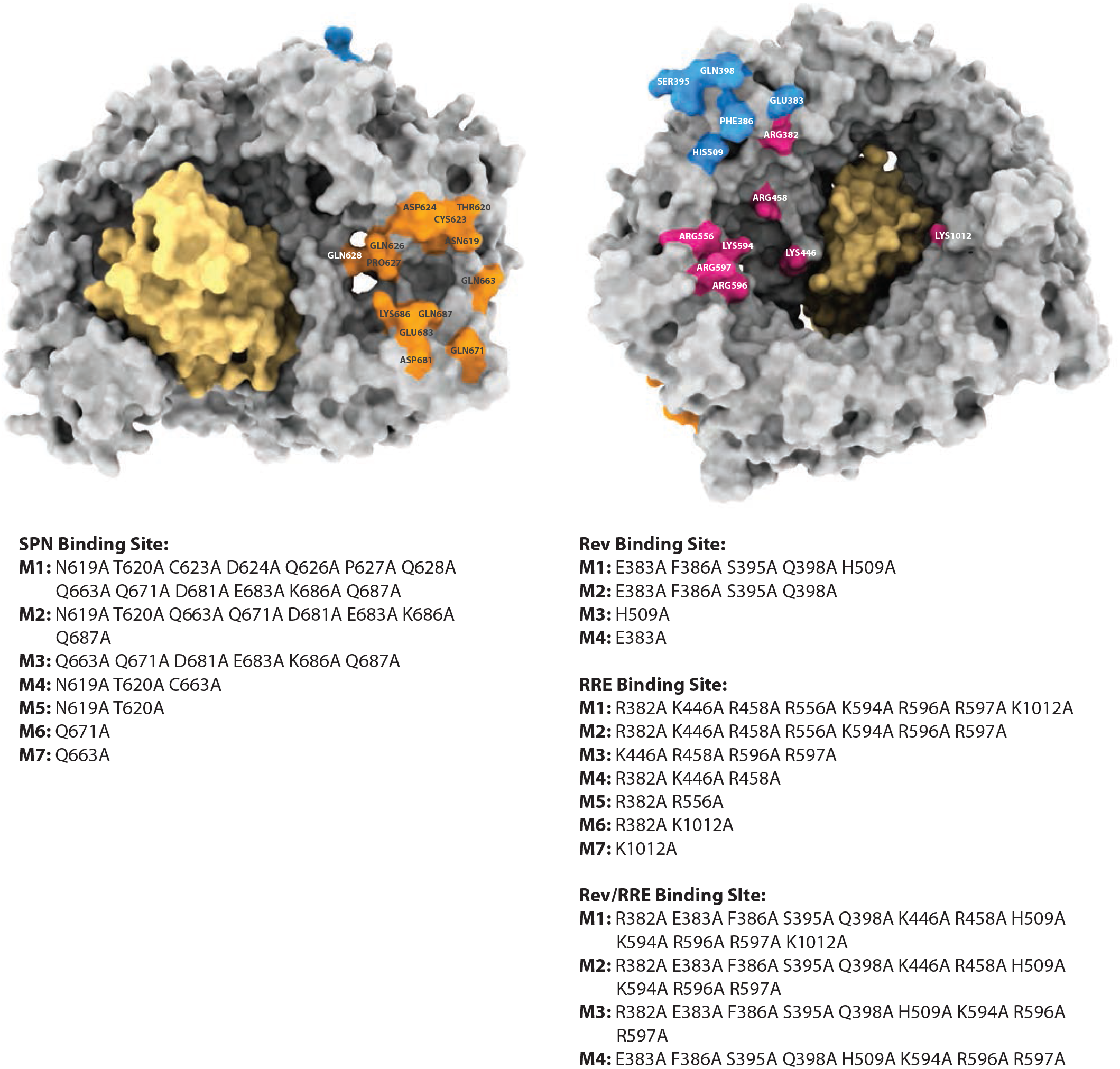
Crm1 Structure Validated Cargo Recognition Sites. Left/Top: Crm1 shown in gray as a surface representation with the SPN binding site residues colored in orange. Left/Bottom: Nomenclature for each construct (M1-M7) and their corresponding mutated residues. Right/Top: Crm1 shown in gray as a surface representation with the Rev and RRE binding site residues colored in blue and pink. Right/Bottom: Nomenclature for each construct (M1-M4 for Rev, M1-M7 for the RRE and M1-M4 for the Rev/RRE) and their corresponding mutated residues.

**Figure S8.**
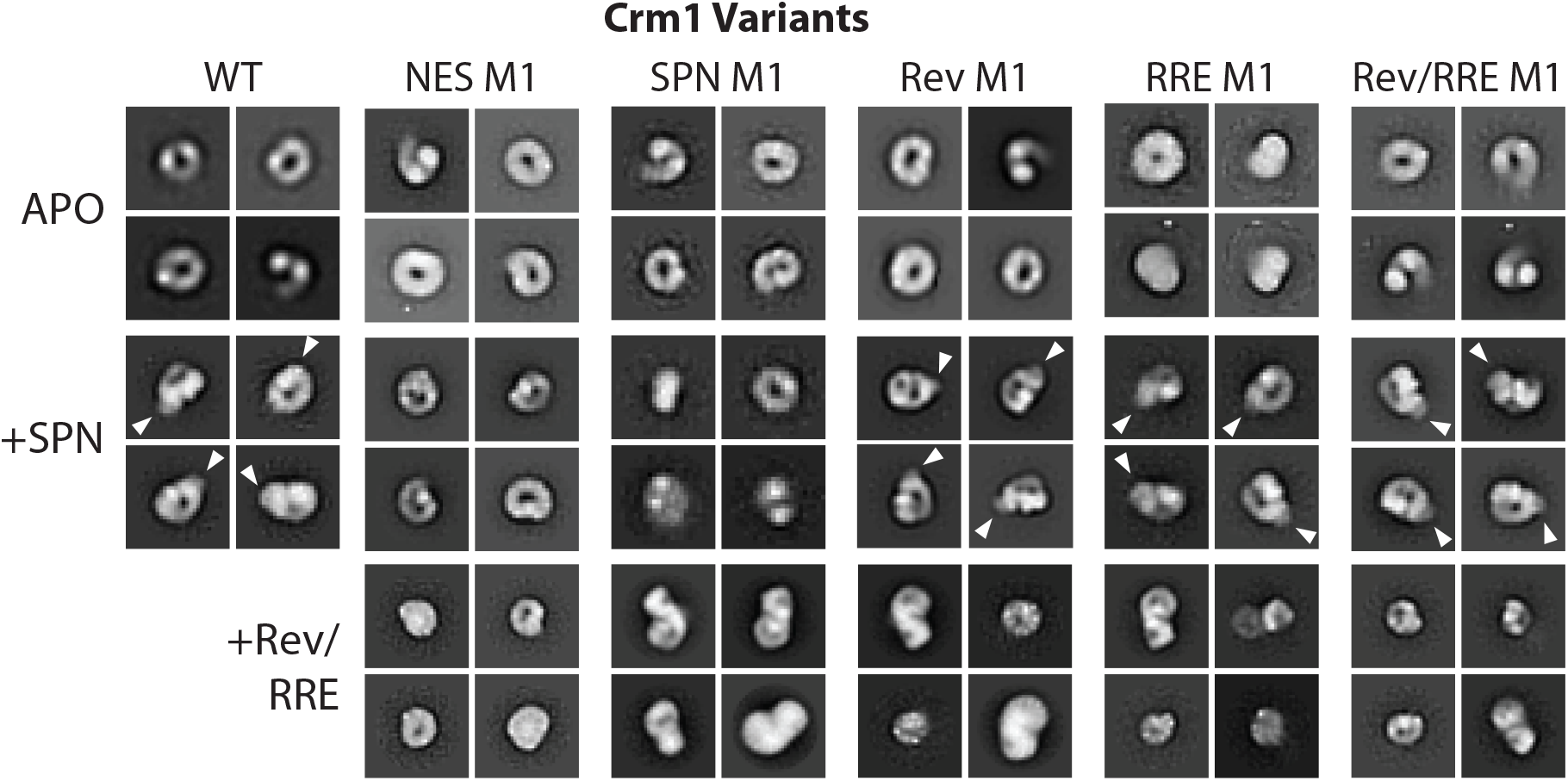
Representative negative stain 2D class averages of Crm1 variants with either Apo, SNP or the Rev/RRE RNP to test efficiency of either overall architecture or complex assembly. Representative 2D class averages of either apo Crm1 (wild type and the M1 construct for each binding site), Crm1/Ran-GTP bound to SPN (same constructs of Crm1 shown as apo), or Crm1/Ran-GTP bound to the HIV-1 Rev/RRE234 RNP (same constructs of Crm1 shown as apo).

**Figure S9:**
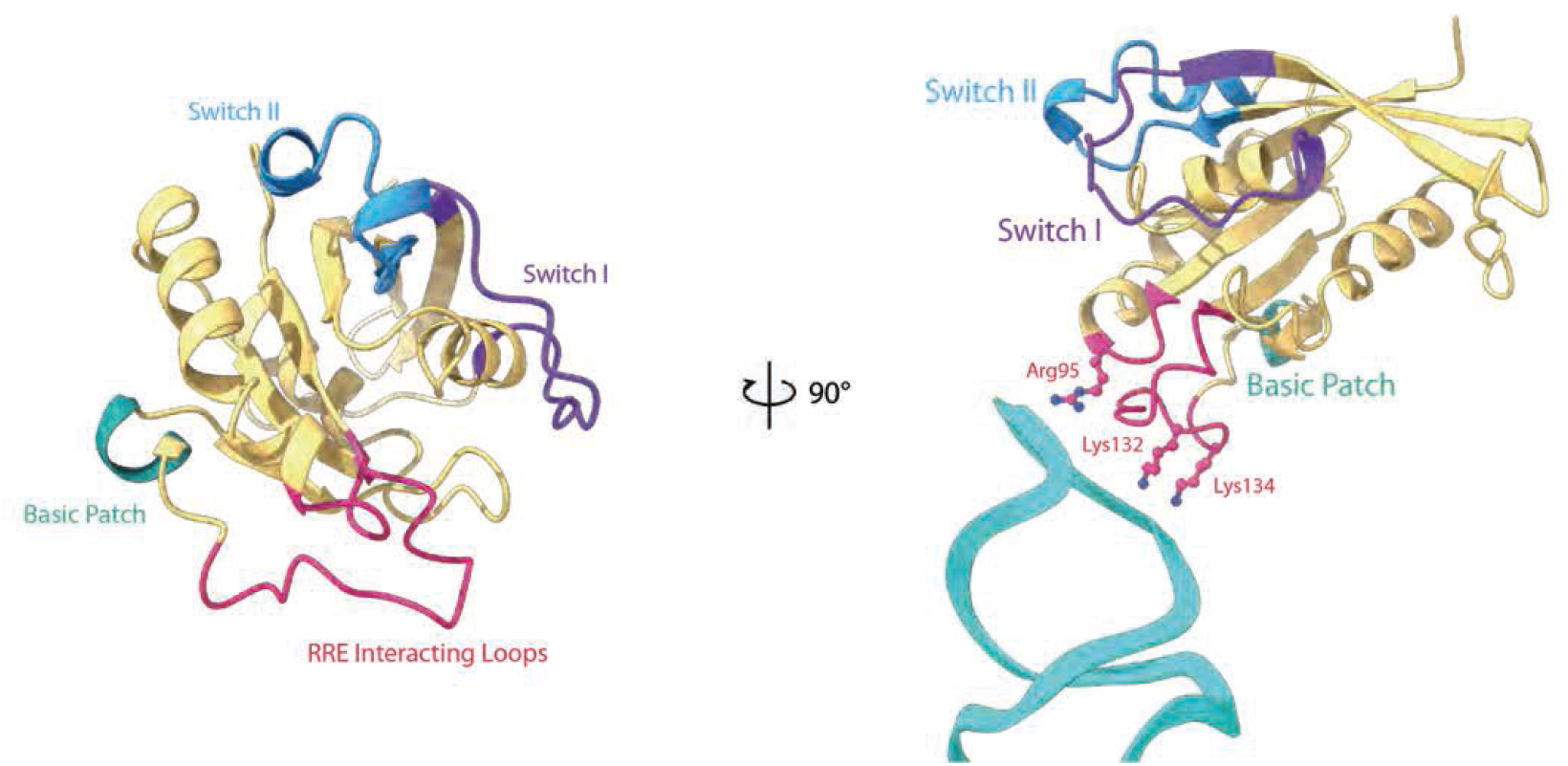
Ran-GTP/RRE interaction. Left: Ran represented in ribbon diagram in peach with switch I and II (responsible for recognizing either GTP or GDP) colored in purple and blue. The basic patch that interacts with Crm1 colored in teal. The two loops that contain the residues utilized by all exportins that traffic RNA and that directly interact with the RRE colored in pink. Right: Ran from the left with the RRE stem IIB represented in ribbon diagram colored in cyan and the interacting residues shown in ball and stick colored in pink.

**Figure S10:**
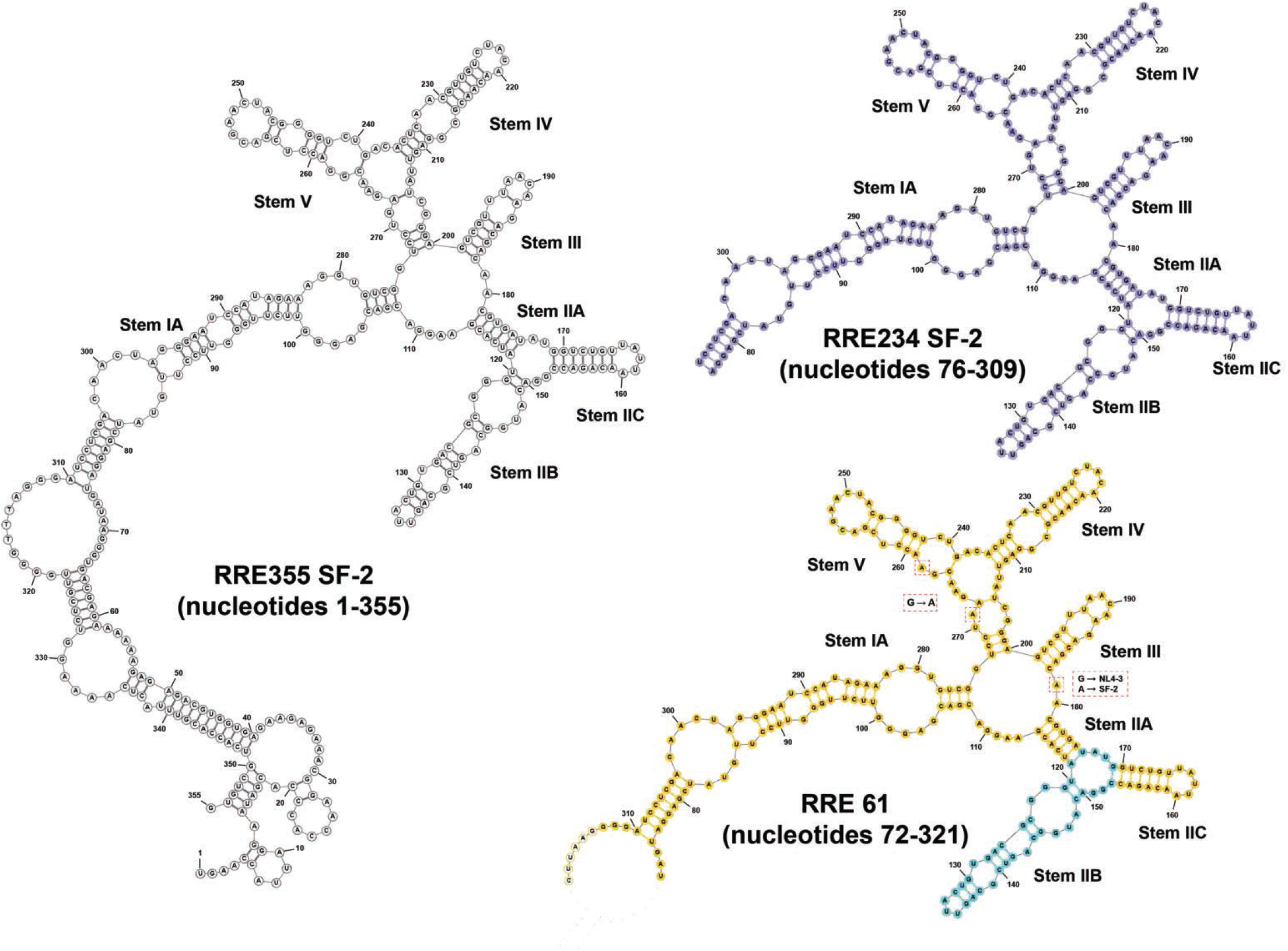
Secondary structure of the HIV-1 SF-2 RRE Variants. Sequence of the full-length HIV-1 RRE355, isolate SF-2, shown in gray. Sequence of RRE 234 (the minimal viable RRE) shown in purple. Sequence of RRE 61 truncated to the minimal RRE, with stem IIB shown in cyan. Mutant G262A and nucleotide A181 are responsible for RRE61 resistance. Position 181 is a guanosine in the NL4-3 isolate but was already an adenosine in the SF-2 isolate. An adenosine at position 269 in the RRE from the NL4-3 isolate is critical to the 3D shape necessary for RRE61 resistance, therefore G269 was mutated to an adenosine in the SF-2 RRE.

**Figure S11:**
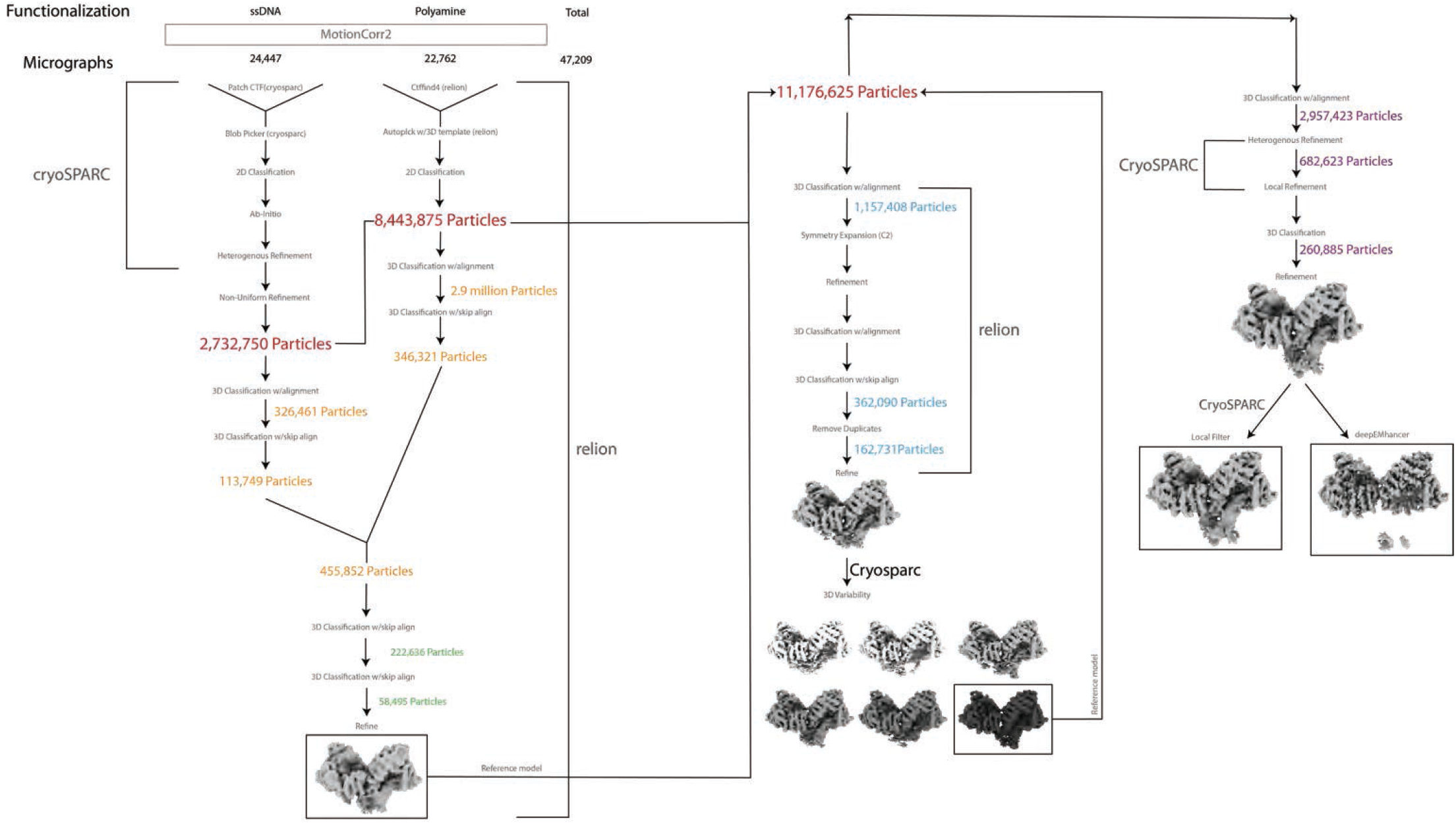
Cryo-EM data processing workflow. Schematic flow-chart representing the image processing approach for the NEC. Thumbnail images of the NEC after refinement at different steps are shown and how each reconstruction was then used as initial reference model to reprocess the initial particle stack to aid in correct particle alignments resulting in an increased overall resolution of the complex.

**Figure S12:**
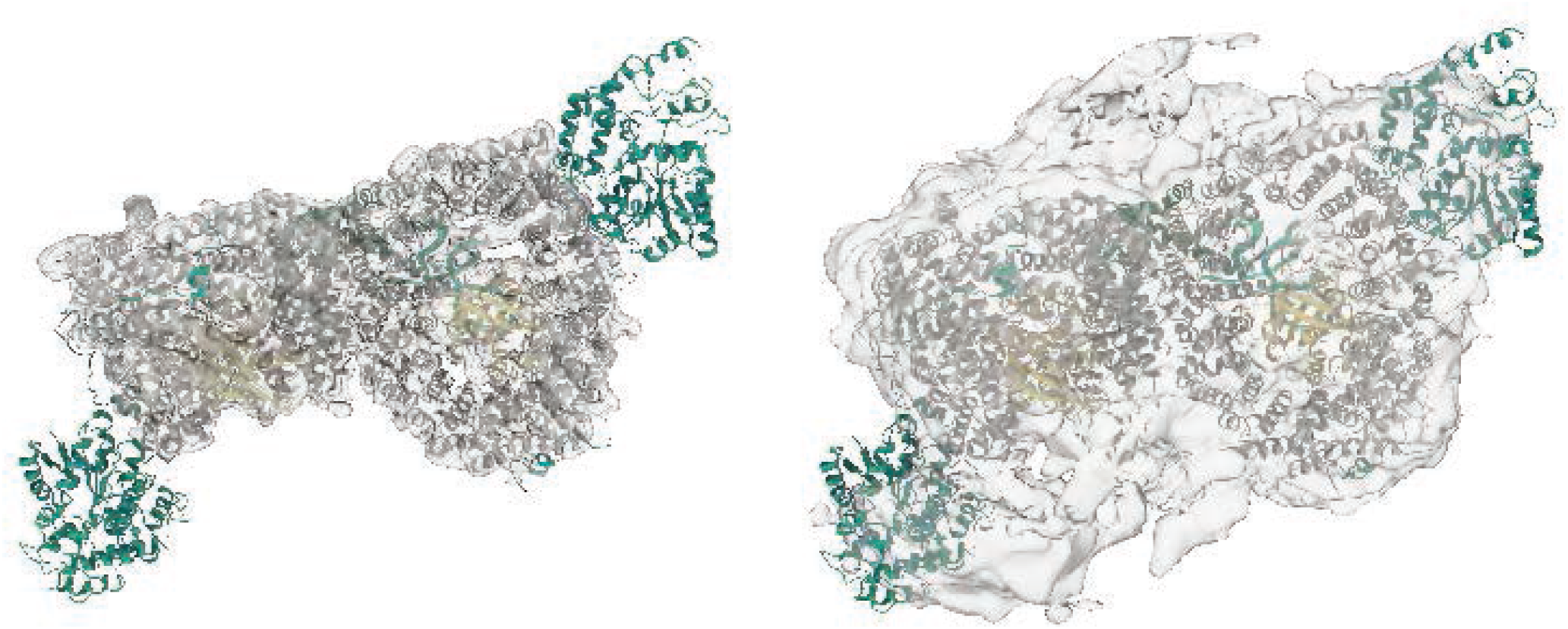
_MBP_Nup214_1916-2033_ Binds to the NEC Complex. Increasing the contour level reveals density for the MBP of the _MBP_Nup214_1916-2033_ construct validating the presence of the Nup214 peptide bound to each Crm1 subunit. Left: The NEC shown at contour levels related to the resolution of the complex. Right: Same map as on the left with increased contour levels.

**Table S1:** Cryo-EM collection, refinement, and validation statistics.

|  | ssDNA oligos | Polyamine |
| --- | --- | --- |
| Data Collection and processing |  |  |
| Microscope | Titan Krios | Titan Krios |
| Magnification | EFTEM 105,000 | EFTEM 105,000 |
| Voltage (kV) | 300 | 300 |
| Electron exposure (e-/Å<sup>2</sup>) | 66 | 68 |
| Defocus range (µm) | -0.8-2 | -0.8-2 |
| Pixel Size (Å) | 0.834 | 0.835 |
| Micrographs collected (no.) | 24447 | 22762 |
| Symmetry imposed | C1 | C1 |
| Initial particle images (no.) | 2,732,750 | 8,443,875 |
| Final Particle images (no.) | 260,885 |  |
| Map resolution (Å) | 3.8 |  |
| FSC threshold | 0.143 |  |
| Map resolution range (Å) | 3.8-142.85 |  |
| Refinement |  |  |
| Initial model used (PDB code) | 5DIS (Crm1/Ran-GTP)/4PMI (Rev dimer with stem IIB mimic) |  |
| Model resolution (Å) | 7.9 |  |
| FSC threshold range (Å) | 0.5 |  |
| Model resolution range | 3.8-142.85 |  |
| Map sharpening B factor (Å) | deepEMhancer |  |
| Model composition |  |  |
| Non-hydrogen atoms | 21,963 |  |
| Protein residues | 2603 |  |
| Nucleotide | 39 |  |
| B factors (Å<sup>2</sup>) |  |  |
| Protein | 325 |  |
| R.m.s deviations |  |  |
| Bond lengths (Å) | 0.003 |  |
| Bond angles (°) | 0.76 |  |
| Validation |  |  |
| MolProbity score | 1.94 |  |
| Clashscore | 18.82 |  |
| Poor Rotamers (%) | 0 |  |
| Ramachandran plot |  |  |
| Favored (%) | 97.03 |  |
| Allowed (%) | 2.89 |  |
| Disallowed (%) | 0.08 |  |

**Table S2:** List of Cnnl residues that contact stem IIB of the RRE.

| Within HEAT Repeats |  |  |
| --- | --- | --- |
| Residue of Crm1 | Crm1 HEAT |  |
| Arg382 | 8B |  |
| Arg458 | 9B |  |
| Lys594 | 12B |  |
| Lys 1012 | 21A |  |
| Within Linkers |  |  |
| Residue of Crm1 | Between HEATS |  |
| Lys446 | 9A | 9B |
| Arg556 | 11B | 12A |
| Arg596 | 12A | 12B |
| Lys597 | 12A | 12B |

**Table S3:**
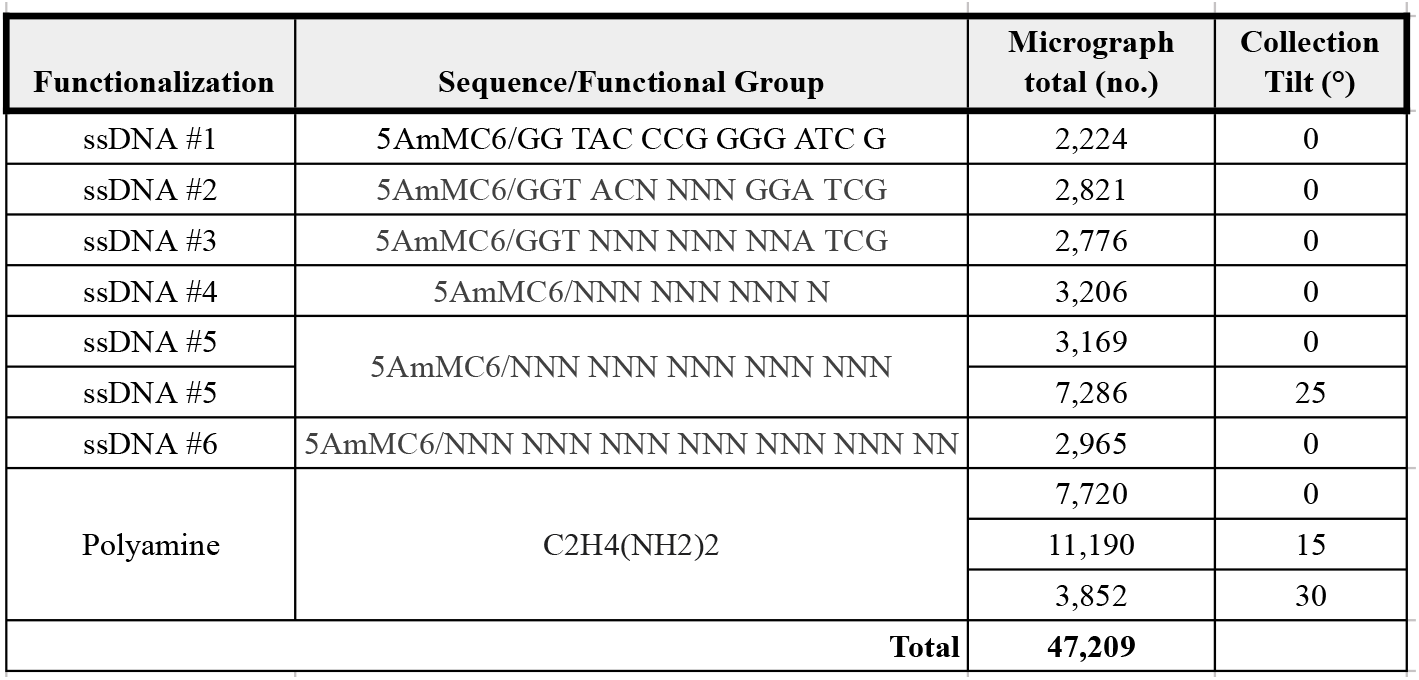
Sequences/functional group, micrograph count and tilt angle for each data set collected.

### Uncropped images of western blots

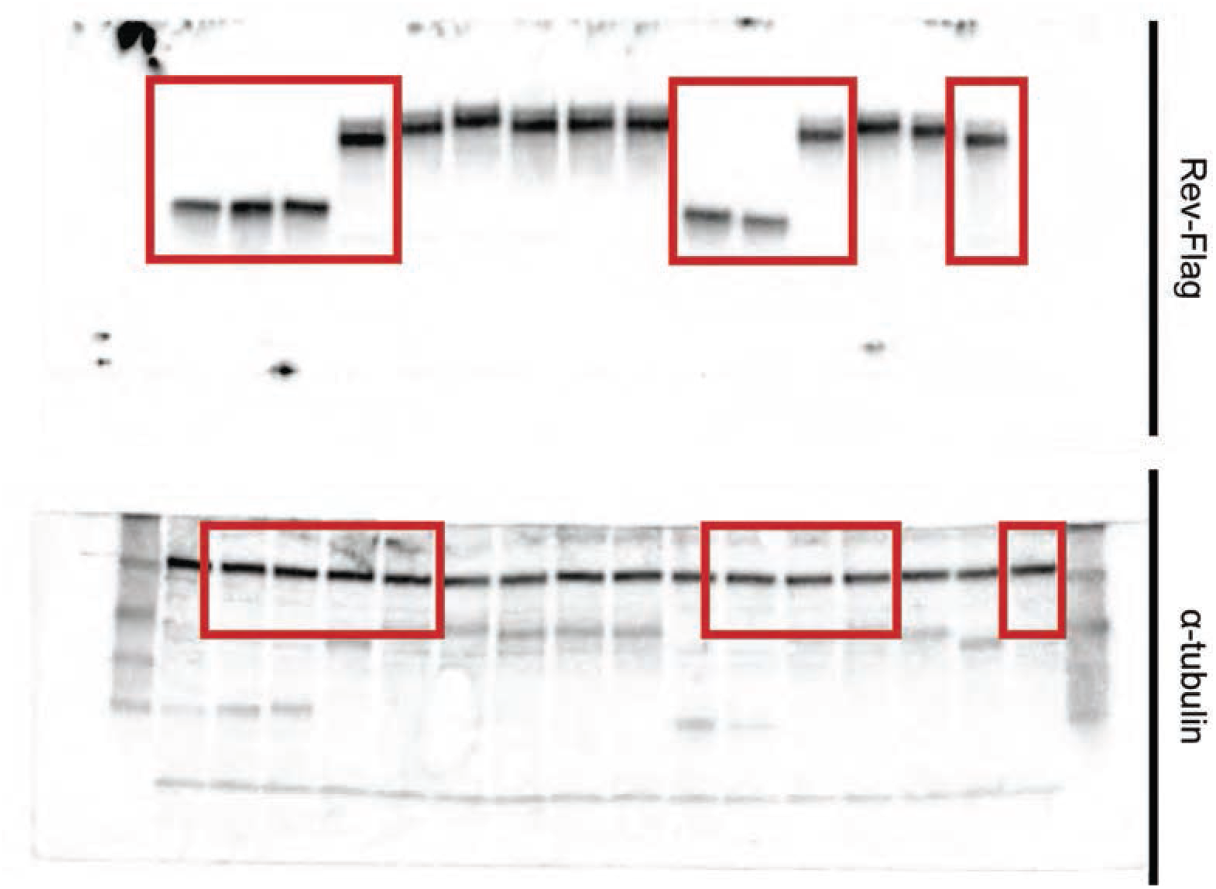

**>Uncropped images of western blot** Immunoblots for Figure 2F. The red rectangles show cropping.

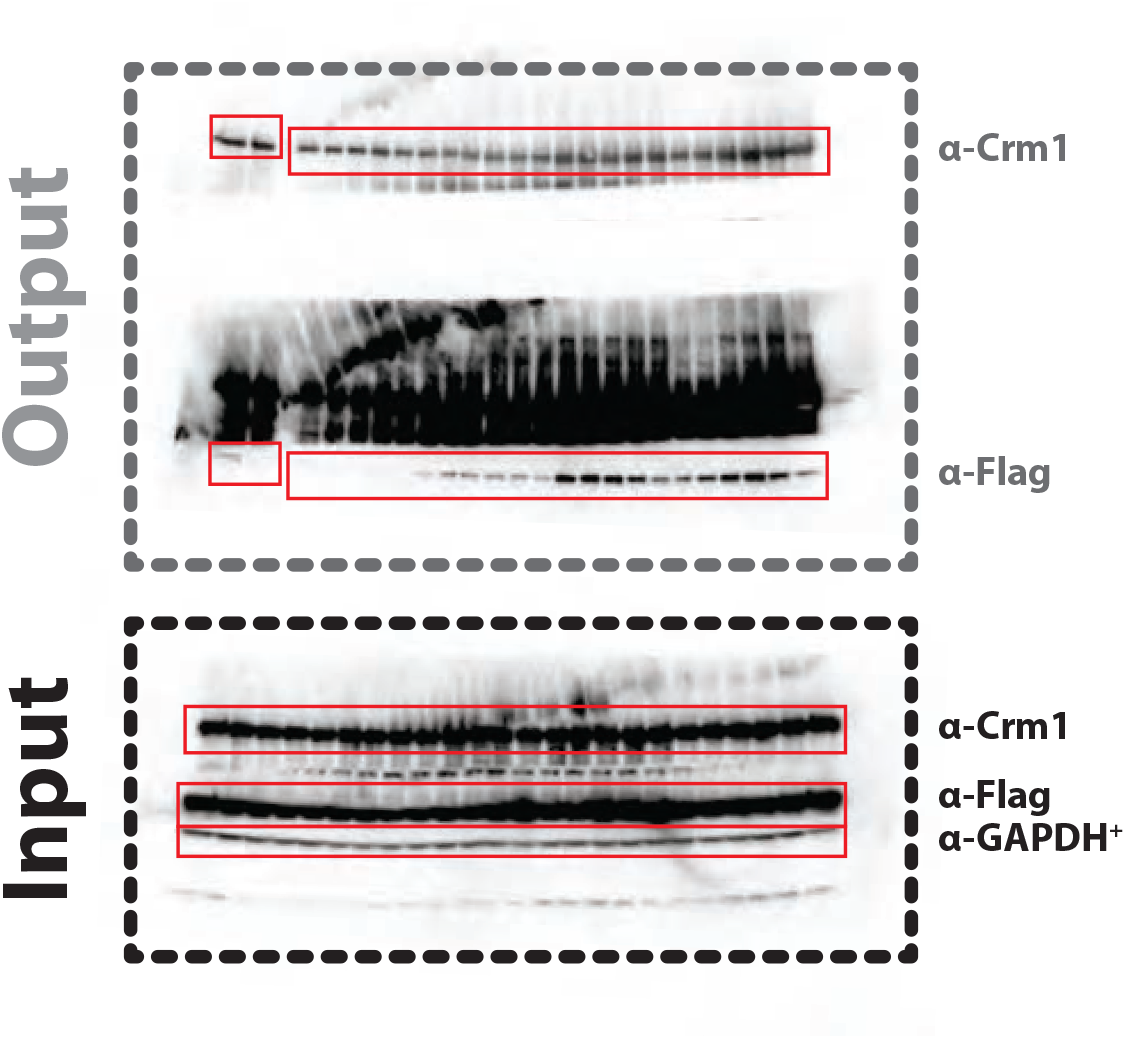

**Uncropped images of western blot**. Immunoblots for Figure 6A. The red rectangles show cropping.

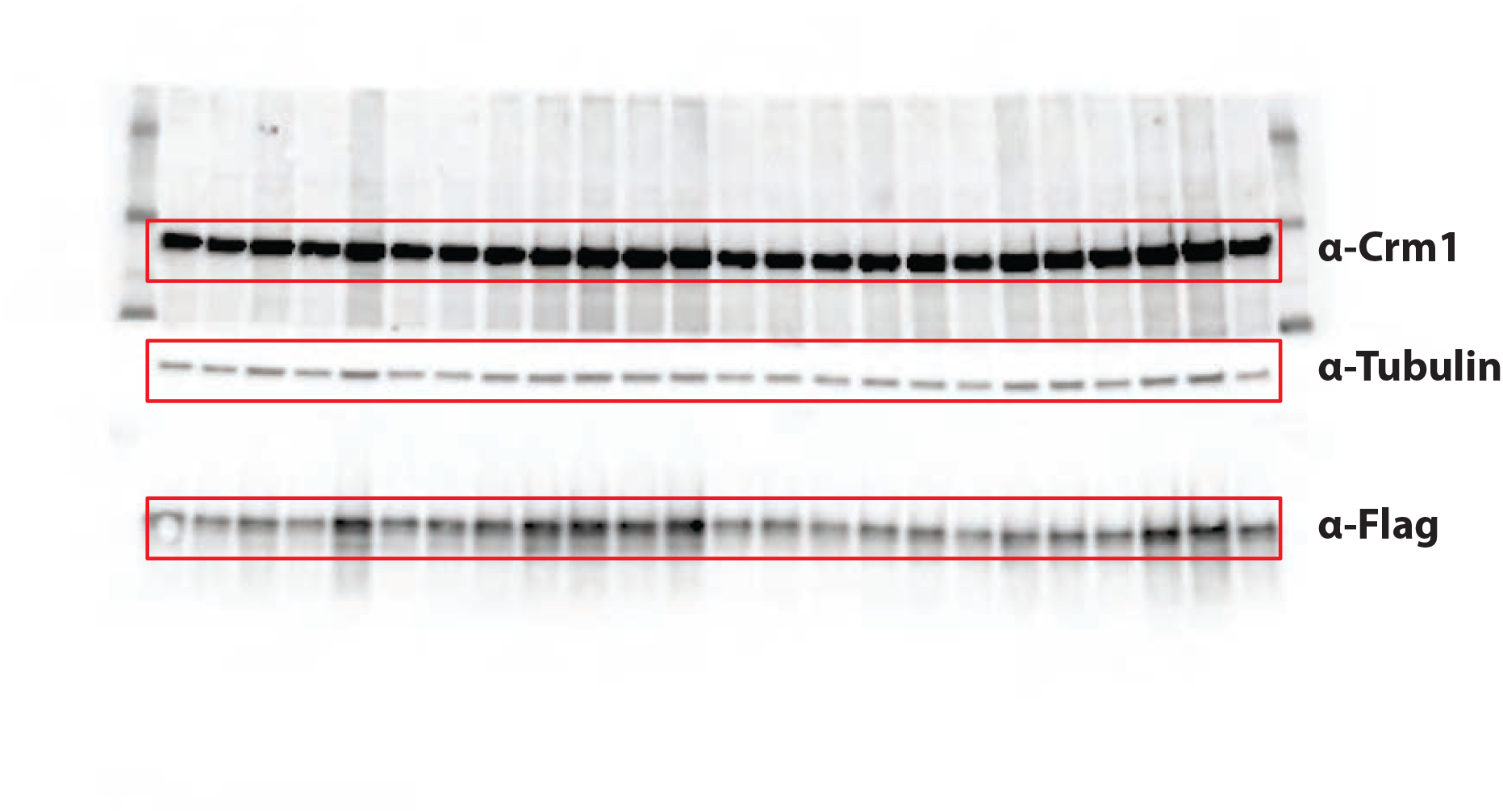

**Uncropped images of western blot**. Immunoblots for Figure 6B. The red rectangles show cropping.

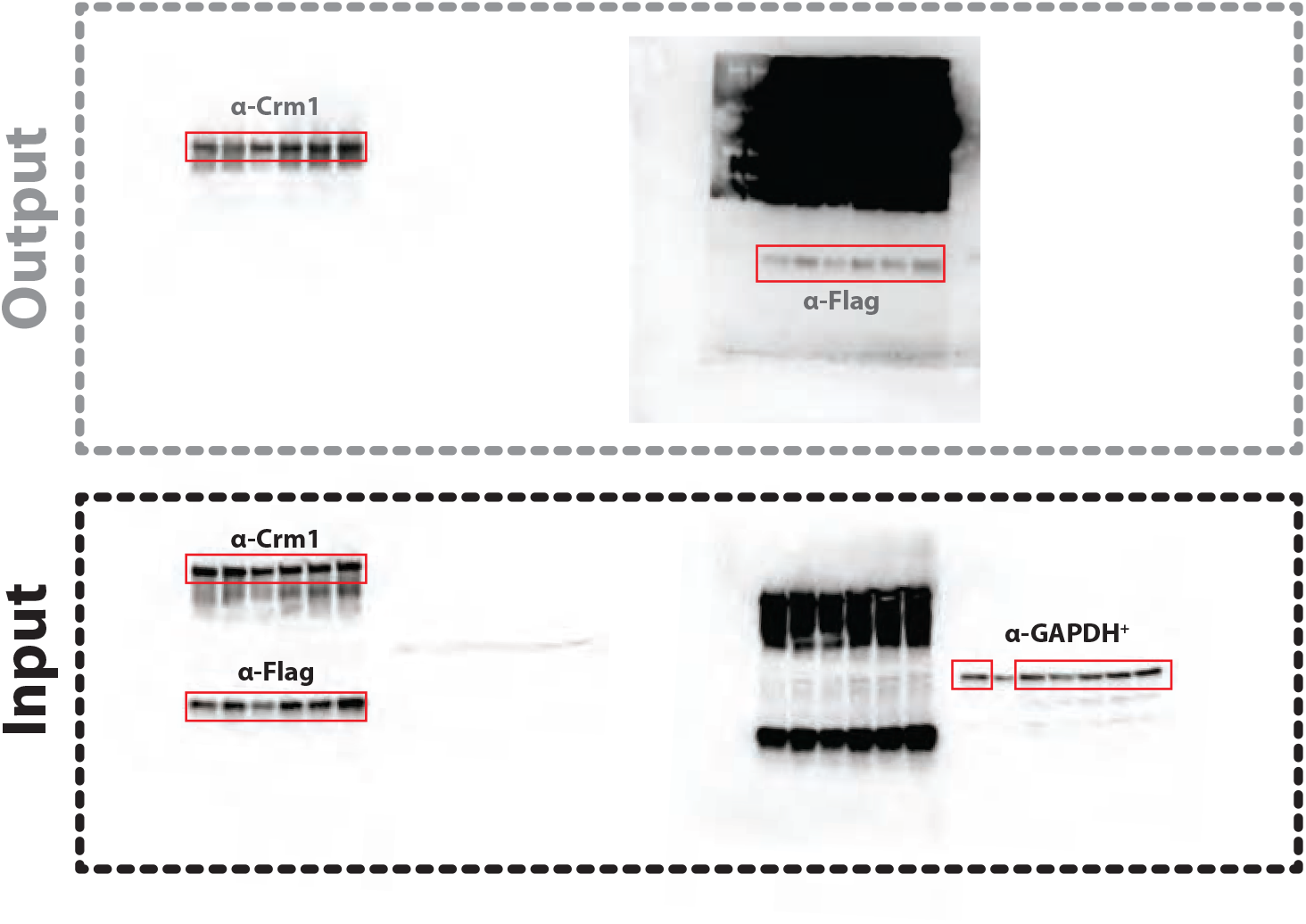

**Uncropped images of western blot. Immunoblots for Figure 6D. The red rectangles show cropping.**

